# Glutamatergic neuron-tumor synapses shape human glioblastoma cell states through radial glia plasticity

**DOI:** 10.64898/2026.05.14.725216

**Authors:** Antoni Martija, Brianna N. Bristow, Dakshesh Rana, Savan Bollu, Elisa Fazzari, Shivani Baisiwala, Claudia V. Nguyen, Weihong Ge, Ryan L. Kan, Daria J. Azizad, Matthew X. Li, Patricia R. Nano, Heejin Cho, Travis Perryman, David A. Nathanson, Kunal S. Patel, Aparna Bhaduri

**Affiliations:** Department of Biological Chemistry, David Geffen School of Medicine, University of California, Los Angeles, California, Los Angeles, CA, USA; Department of Cellular and Physiological Sciences, Faculty of Medicine, Life Sciences Institute, University of British Columbia (UBC), Vancouver, BC, Canada; Department of Neurosurgery, David Geffen School of Medicine, University of California, Los Angeles, California, Los Angeles, CA, USA; Department of Molecular and Medical Pharmacology, David Geffen School of Medicine, University of California, Los Angeles, Los Angeles, CA, USA

## Abstract

Glioblastoma (GBM) is a devastating primary brain tumor with remarkable inter- and intra-tumoral heterogeneity. GBM cells assume a spectrum of neurodevelopmental-like phenotypes and co-opt normal neurophysiological processes, which include synaptic integration with their neuronal microenvironment. This is mediated by neuron-tumor synapses (NTS) that predominantly involve glutamatergic receptors, which drive calcium elevations that promote tumor proliferation and invasion. The exact relationship between synaptic signaling and tumor cell fate specification, however, remains largely unexplored. Here, we develop and leverage a synapse-optimized human organoid tumor transplantation (so-HOTT) model of GBM to decipher how glutamatergic signaling impacts GBM lineage trajectories. so-HOTT preserves patient tumor heterogeneity, features excitatory NTS, and enables clonal lineage tracing of tumor cells after NTS perturbations. Genetic and pharmacological inhibition of AMPA and kainate receptors in so-HOTT shifts tumor cell composition from neuronal fates toward progenitor-proximal astrocytic/mesenchymal states. This occurs through the attenuation of calcium signaling and reduced plasticity of malignant radial glia (RG)-like progenitors, a previously unrecognized target of NTS. Through the integration of inputs from the neuronal microenvironment into glutamatergic signaling, progenitor populations modulate their transcriptional programs and cell fate, ultimately shaping GBM tumor heterogeneity. Targeting synaptic input may thus constrain the heterogeneity that fuels GBM adaptation and therapeutic escape.

## Introduction

Functional integration of glioblastoma (GBM) in neural circuits through neuron-tumor synapses (NTS) has emerged as a critical mechanism driving tumor progression^1–5^. Landmark studies in cancer neuroscience have shown that in animal models, human organotypic slice cultures, and neuron-glioma 2D co-culture systems, these NTS transmit excitatory signals that elicit calcium activity, which then promotes tumor proliferation and/or invasion^1–7^. To communicate with their neuronal microenvironment, tumor cells from GBM and other glioma subtypes predominantly utilize glutamatergic ⍺-amino-3-hydroxy-5-methyl-4-isoxazole propionic acid (AMPA) receptors (AMPARs)^1,2,6,7^; however, in recent years, other neurotransmitters and neurotransmitter receptors (NTRs) have also been implicated^3,4,8,9^. Interestingly, the synaptogenic potential of GBM cells is enriched in tumor cell states with neurodevelopmental signatures, which are coincidentally enriched at the invasive tumor margin^4,10–12^. This suggests that the neuronal tumor microenvironment can actively shape tumor cell identities to become more neurodevelopmental-like. In support of this, previous work in the field of neurodevelopment has similarly shown that neural progenitor cells (NPCs) express various NTRs including AMPARs, and that perturbing these receptors influences the cell fate trajectories of these progenitors^13–18^. While numerous studies in the field of cancer neuroscience have already solidified the role of synaptic input on tumor proliferation and invasion, a direct functional link between NTS and GBM cell fate specification has yet to be fully explored. Investigating this link is important, given that inter- and intra-tumoral heterogeneity is immensely pervasive in GBM^19–24^ and is thought to increase the potential of tumor cell adaptability to therapeutic pressures and a continuously evolving microenvironment^25,26^. We hypothesize that in GBM, excitatory synaptic signaling through glutamatergic receptors not only promotes classical cancer hallmarks of proliferation and invasion, but also contributes to GBM heterogeneity by actively shaping GBM plasticity and the resulting cell states.

Investigating the impact of NTS on GBM cell fate trajectories requires a human-specific model that 1) harbors these synapses, 2) allows the tractable and scalable modulation of neuron-tumor interactions, and 3) permits interrogations on tumor cell types and their lineage relationships. The provision of a human-specific context is critical, given the known evolutionary differences between mice and humans in terms of neuronal and glial populations^27–29^ and species-specific differences in the nature of synaptic interactions^30,31^. Additionally, permissive effect of a 3D cytoarchitecture on GBM cell type preservation and invasion^32–37^ benefits from a complex neuronal microenvironment. Existing human organotypic slice cultures already address these limitations^5,38^; however, these models are difficult to acquire and are less amenable to independent perturbations of tumor and microenvironmental cells. To address this, various human cortical organoid models of GBM have emerged, including genetically transformed cerebral organoids^34,39^, GBM tumor explants^40^ and assembloids generated by the transplantation of glioma stem cells or explants into cortical or cerebral organoid scaffolds^35,36,41^.

We developed a human organoid tumor transplantation (HOTT) model of GBM, which features fresh, surgically resected primary tumor cells engrafted into stem cell-derived cortical organoids^37^. This approach preserves the heterogeneity of GBM cell types, while allowing organoid-specific perturbations^37^, modular addition of other relevant microenvironmental cell types like immune cells^42^, or modification of the neural organoid scaffold identity^43^. Importantly, this model has been successfully used to map lineage relationships and clonal dynamics of diverse GBM cell types at baseline and in response to therapy^24,44^. Despite these critical advances, current assembloid models still lack substantial NTS formation, owing to issues like developmental immaturity of the cortical organoid scaffold and suboptimal culture conditions.

To overcome these limitations, we built upon HOTT to develop a synapse-optimized human organoid tumor transplantation (so-HOTT) model of GBM. We validated the fidelity of so-HOTT to human patient tumor samples, established the functionality of the synapses generated, performed genetic and pharmacological perturbations to dissect the impact of synaptic signaling on tumor cell states, and applied lineage tracing to discover how glutamatergic signaling through radial glia-like cells, a neurodevelopmental cell type reactivated in GBM, modulates GBM plasticity. These findings establish progenitor-specific synaptic input as a critical determinant of GBM evolution and a potential therapeutic target for constraining the heterogeneity of these complex brain tumors.

## Results

### so-HOTT Induces Synaptic Gene Expression and Intercellular Interactions

To develop a system in which we can study human NTS, we built upon our previous innovations in the HOTT model^37^. Freshly resected core and peripheral regions of primary tumors, infected with EGFP lentiviruses for 90 mins after dissociation, were transplanted into pre-grown Week 10-12 cortical organoids using our previously described hanging drop method^37^ (**Fig. 1A-B**). We utilized the same media conditions as HOTT for our control; for so-HOTT, we further supplemented the HOTT media with B-27 with Vitamin A and doubled the amount of Matrigel to support neuronal viability. Additionally, we administered rationally selected maturation factors to promote NTS formation in so-HOTT. The small molecules and growth factors added to so-HOTT included brain-derived neurotrophic factor (BDNF), which has been shown to promote AMPA receptor membrane trafficking and glutamatergic currents in glioma^7^; neuroligin-3 (NLGN3), which increases synaptic gene expression in tumor cells^45^; and the four-part GENtoniK cocktail, which has been effective in increasing synapses within cortical organoids alone^46^. Two components of this cocktail (Bay K 8644 and N-methyl-D-aspartate or NMDA) promote calcium-dependent transcription, while the other two (GSK2879552 and EPZ-5676) remodel chromatin to increase the expression of synaptic genes and ion channel subunits^46^.

**Figure 1.**
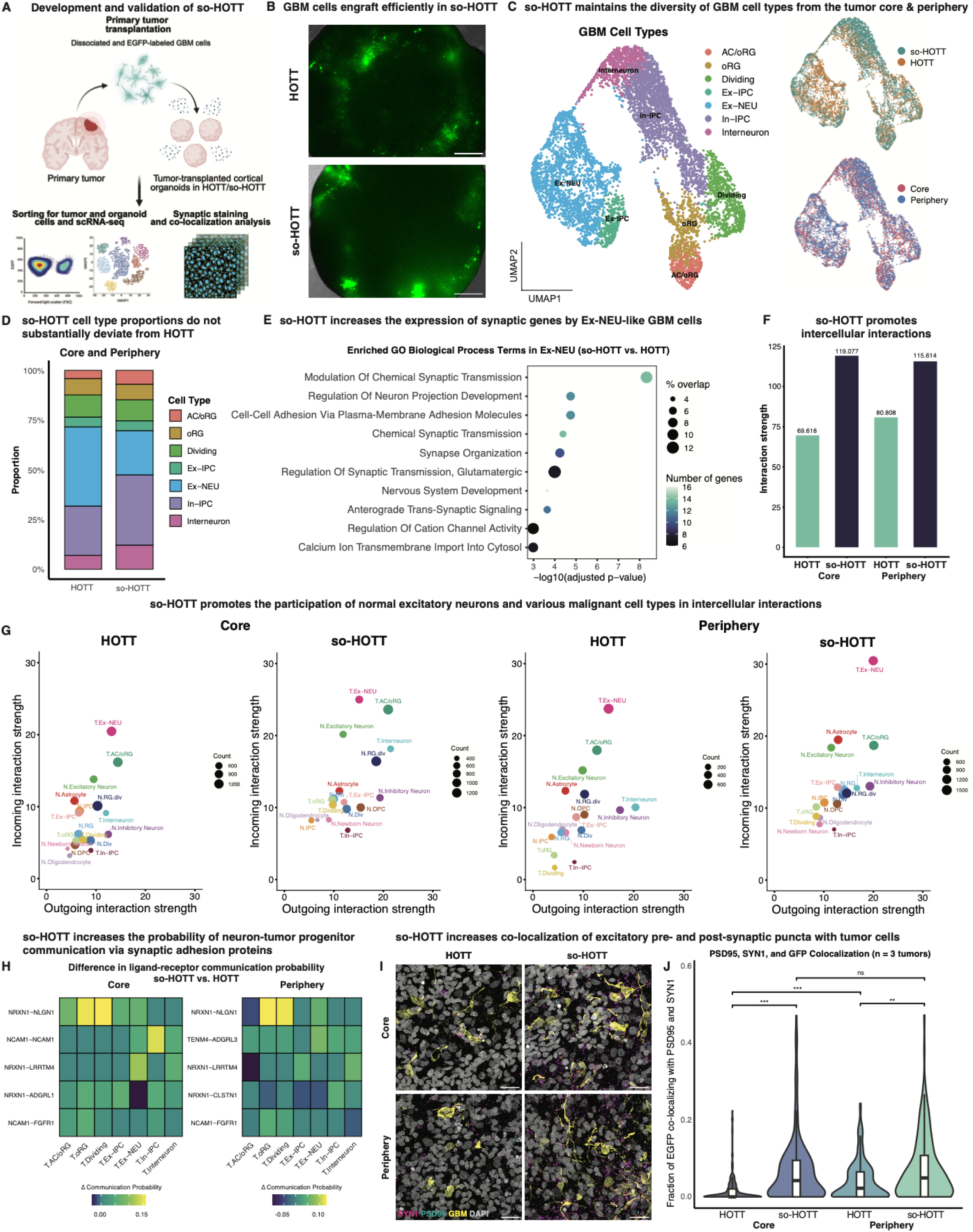
so-HOTT Induces Synaptic Gene Expression and Intercellular Interactions A) To optimize a system that can recapitulate neuron-tumor synapses (NTS), we leveraged the existing Human Organoid Tumor Transplantation (HOTT) system and made modifications aimed at promoting neuronal maturation and synaptic gene expression. Schematic shows how primary tumor cells (directly from surgical resection, never cultured) are dissociated, labelled with lentiviral EGFP and transplanted into a pre-existing Week 10-12 cortical organoid that mimics the tumor microenvironment. Experiments were performed using glioblastoma (GBM) cells from the tumor core and tumor periphery. Using different media conditions (HOTT and so-HOTT), we tested if a cocktail of supplements (Methods) could drive NTS formation. To validate the efficacy of so-HOTT, we sorted for both EGFP+ tumor cells and EGFP- organoid cells and performed single-cell RNA-sequencing (scRNA-seq). Synaptic stains were also performed to validate NTS formation at the protein level. B) Live images of HOTT and so-HOTT organoids 2 weeks post-transplantation show the effective engraftment of primary patient tumor cells (EGFP+) in cortical organoids. Scale bar = 500 µm C) Uniform Manifold Approximation and Projection (UMAP) of scRNA-seq after cell types are mapped to a reference atlas of GBM and annotated based upon marker gene analysis shows heterogeneity of cell types as expected in HOTT and so-HOTT (left UMAP). Major cell types were represented in both HOTT and so-HOTT (right, top) conditions and across core and peripheral samples (right, bottom). D) Stacked bar plot shows cell type proportions with limited changes in cell type heterogeneity in the so-HOTT condition compared to HOTT. E) Gene Ontology (GO) Biological Process Term enrichment of the excitatory neuron-like tumor cells (EX-NEU) shows a significant increase of programs related to synaptic transmission and neuronal projection in the so-HOTT system. The x-axis shows statistical significance (-log10 of adjusted p-value) of the GO terms listed on the y-axis. Color scale corresponds to the number of genes for each term, while dot size shows the percentage of genes overlapping with the total number of genes for each term. F) Bar plot showing predicted cell-cell interaction strengths using CellChat. Across core and periphery tumor samples, the so-HOTT condition, which we call synapse-optimized HOTT (so-HOTT), had more intercellular interactions overall compared to HOTT. G) Dot plots show CellChat-predicted outgoing interaction strength (x-axis) and incoming interaction strength (y-axis) with dots scaled to the number of inferred links. Interaction strength is an arbitrary unit. Detailed examination of the cell types involved in predicted intercellular interactions shows that so-HOTT drives intercellular communication between non-malignant neurons and a variety of tumor cell types, most notably Ex-NEU-like and AC/oRG-like cells. H) Heatmaps showing the top differential ligand-receptor (L-R) pairs utilized by excitatory neurons (sender) to communicate with various tumor cell types (receiver) across so-HOTT and HOTT. Tumor cell types are listed on the x-axis, while L-R pairs are on the y-axis. Color scale corresponds to the difference in L-R communication probability, which can range from 0.0 to 1.0, between so-HOTT and HOTT. The largest enrichment was observed for progenitor populations NRXN1-NLGN1. I) Immunostainings showing the expression of presynaptic marker SYN1 (magenta), excitatory postsynaptic marker PSD95 (cyan), and EGFP (yellow) marking GBM cells across core and peripheral regions and HOTT/so-HOTT media conditions. Nuclei were stained with DAPI (gray). Representative images were from tumor LB5658 transplanted in cortical organoids. Scale bar = 20 µm. J) Quantification of presynaptic and postsynaptic puncta colocalization with EGFP was performed to identify the fraction of tumor signal co-localizing with PSD95 and SYN1. For both tumor core and periphery, so-HOTT had significantly more excitatory synaptic interactions than HOTT. Data were analyzed using a Kruskal–Wallis test followed by Dunn’s post-hoc test with Holm correction for multiple comparisons. Significance is indicated as follows: *, p ≤ 0.05; **, p ≤ 0.01; ***, p ≤ 0.001; ns, not significant.

To compare the resulting HOTT and so-HOTT models, we performed single-cell RNA-sequencing (scRNA-seq) on both EGFP+ tumor cells as well as the EGFP- cells from the organoid microenvironment (**Fig. 1B**) after a four-week of organoid maintenance (Methods). After using stringent quality control (QC) metrics to identify high-quality malignant single cells for downstream analysis (Methods), cell types were annotated by cluster marker annotation and comparison to existing GBM atlases^19,24^ (**Fig. 1C, SFig. 1A-C**). We observed that with so-HOTT, we can still preserve the diversity of cell types similar to our standard HOTT model (**Fig. 1D**, **SFig. 1A**). Examination of differentially expressed genes (DEGs) globally revealed upregulation of RNA biosynthesis, neuronal differentiation, and cell migration in so-HOTT (**SFig. 1D**). DEG analysis in a cell-type-specific manner highlighted maximal changes in the neuronal-like population, with limited shifts in progenitor-like tumor cells such as excitatory intermediate progenitor-like (Ex-IPC) and astrocyte/outer radial glia-like cells (AC/oRG) (**SFig. 1F**). Gene Ontology (GO) analysis of the DEGs in excitatory neuron-like (Ex-NEU) cells revealed enrichment of synaptic transmission and glutamatergic signaling (**Fig. 1E**), consistent with known engagement of this tumor cell type in synaptic interactions^2^. Thus, so-HOTT induces synaptogenic gene programs without compromising tumor heterogeneity.

Given several GO terms associated with synaptic communication and cell-cell adhesion, we sought to investigate how interactions between the tumor and the microenvironment were changing with so-HOTT. To explore this, we leveraged CellChat-based cell-cell communication analysis^47^, finding that overall interaction strength between cells was increased in so-HOTT (**Fig. 1F**). We looked at the cell types participating in these intercellular interactions and found greater outgoing interaction strength of non-malignant excitatory neurons from the organoid microenvironment, as well as increased incoming interaction strengths of Ex-NEU- and AC/oRG-like tumor cells when grown in so-HOTT (**Fig. 1G**). Thus, so-HOTT increases the ability of normal excitatory neurons to act as signaling senders, while promoting the ability of certain tumor cell types to receive incoming signals. Interestingly, within each media condition, Ex-NEU-like tumor cells from the tumor periphery had higher interaction strengths compared to their equivalents from the core region, suggesting that so-HOTT preserves known intrinsic differences between cells from the tumor core and the invasive margin^4,10–12^. so-HOTT peripheral tumor cells also exhibited enrichment of genes related to oxidative phosphorylation (**SFig. 1E**), consistent with their known preference for mitochondrial respiration over glycolysis, the latter being more pervasive in hypoxic core regions^11,48^.

Previous investigations of glioma cell interactions with the neuronal microenvironment have focused on OPC-like and NPC-like populations^2,6^. Interactions involving neuron-like tumor cells have been described^2^, but this specific population has not been well-characterized. This is likely because malignant neuron-like populations distinct from NPC-like cells were only recently characterized transcriptomically through GBM atlases^20,24^. Our system enables investigation of this cell type, as well as malignant progenitors and their engagement with the neuronal microenvironment. To determine how these various tumor cell types communicate with excitatory neurons in their environment, we performed ligand-receptor analysis using CellChat. We found that synaptic pathways^49^, including NRXN, ADGRL, and CADM, were the top mediators of neuron-tumor communication in so-HOTT (**SFig. 1G)**. Core and peripheral tumor cells equally utilized these pathways (**SFig. 1H**), although distinct differences that likely reflect regional biases in interaction modes were also observed. Among ligand-receptor pairs mediating these synaptic interactions, NRXN1-NLGN1 were found to be the most enriched in so-HOTT compared to HOTT, consistent with NTS development (**Fig. 1H)**. These enriched interactions mostly involved oRG-like and dividing tumor cells, indicating that so-HOTT also facilitates progenitor involvement in excitatory synaptic interactions.

To confirm if the synaptic interactions predicted by CellChat were indeed increased in so-HOTT, we performed immunostainings of presynaptic (SYN1) and postsynaptic (PSD95) markers, measuring the colocalization of these puncta with EGFP-labeled core and peripheral tumor cells transplanted in HOTT or so-HOTT systems. We observed synaptic puncta at the junction of EGFP-positive patient tumor cells and microenvironmental organoid cells across these conditions. As predicted from the scRNA-seq data, there was an increase in the number of SYN1+ PSD95+ excitatory synapses in the so-HOTT system compared to HOTT across core and peripheral tumor cells (**Fig. 1I-J**). Consistent with the predicted baseline differences in synaptic engagement of core and peripheral tumor cells, we also observed a significantly higher colocalization ratio in peripheral tumor cells in standard HOTT conditions (**Fig. 1J**). However, this difference was attenuated in so-HOTT, indicating that the optimized media coaxes core and peripheral GBM cells to form excitatory NTS at almost equal rates. Taken together, our so-HOTT system is able to simultaneously preserve key identities and features of GBM cells and promote their interactions with the neuronal microenvironment provided by the cortical organoid scaffold.

### so-HOTT Mimics Synaptic and Functional Features of Primary Patient GBMs

Studies thus far have highlighted that tumors are primarily postsynaptic^1,3,4,6^ and we sought to validate if this pattern persists in so-HOTT. We observed that presynaptic and postsynaptic Synaptic Gene Ontologies (SynGO) module expression^50^ is comparable between atlas datasets of primary patient GBM^24^ and so-HOTT. Importantly, in both cases, the expression of postsynaptic modules was significantly higher than presynaptic modules, consistent with expected ratios (**Fig. 2A**).

**Figure 2.**
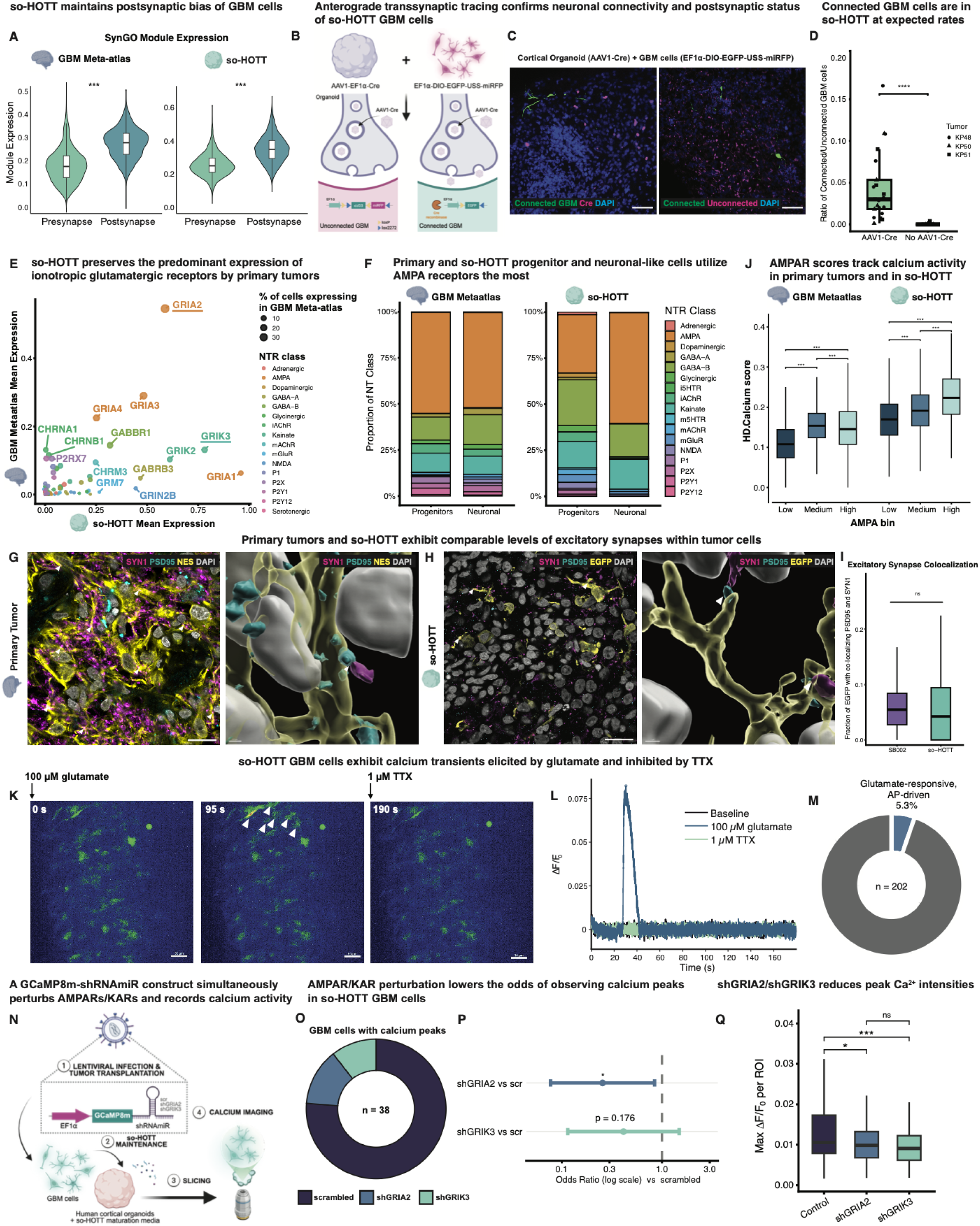
so-HOTT Mimics Synaptic and Functional Features of Primary Patient GBMs A) To determine if so-HOTT preserves the known postsynaptic status of primary GBM cells, we scored tumor cells from the GBM Meta-atlas ^24^ and so-HOTT for genes that are known to be exclusively presynaptic or exclusively postsynaptic based on SynGO ^50^. Primary tumor cells show higher expression of SynGO of postsynaptic than presynaptic genes, as expected, and this was recapitulated in so-HOTT GBM cells. Pairwise comparisons were performed using two-sided unpaired Wilcoxon rank-sum tests (***p ≤ 0.001). B) To functionally validate that so-HOTT GBM cells are indeed postsynaptically connected to neurons in the organoid microenvironment, we performed anterograde transsynaptic tracing using an AAV1-Cre virus ^51,52^ introduced into a subset of cortical organoids prior to tumor transplantation. Tumor cells harboring a Cre-dependent, double-floxed inverse orientation (DIO) EGFP reporter under the control of an EF1⍺ promoter were then transplanted into organoids with or without AAV1-Cre. This reporter also contains an miRFP670 cassette that is excised upon Cre recombination. By default, unconnected tumor cells will be exclusively miRFP670-positive. If a tumor cell is postsynaptically connected to a neuron transduced by AAV1-Cre, the said tumor cell will be expected to express EGFP instead of miRFP670 after AAV1-Cre transsynaptic spread and Cre-dependent recombination. C) Immunostainings showing EGFP+ connected GBM cells in the vicinity of Cre-positive neurons in organoids pre-transduced with AAV1-Cre (left). An alternate immunostaining panel shows EGFP+ connected cells amidst miRFP670+ unconnected cells (right). The experiment was performed across n = 3 tumors. Scale bar = 50 µm. D) Quantification of the ratios of connected/unconnected GBM cells (y-axis) in the AAV1-Cre and negative control conditions (x-axis) across n=3 tumors indicates that about 5% of tumor cells are synaptically connected in so-HOTT. Pairwise comparisons were performed using two-sided unpaired Wilcoxon rank-sum tests (****p ≤ 0.0001). E) Scatter dot plot showing the mean expression of various neurotransmitter receptors (NTRs) in so-HOTT (x-axis) versus in primary tumors from the GBM Meta-atlas (y-axis). Each receptor is color-coded by its receptor family, which was assigned based on the neurotransmitter it binds to and whether it acts through an ionotropic (ligand-gated) or metabotropic (GPCR-coupled) manner. Dot sizes correspond to the percentage of primary tumor cells expressing a particular NTR. The comparison shows that AMPA receptors, particularly GRIA2, are highly expressed in both the primary tumor context and in so-HOTT. Kainate (GRIK3, GRIK2), GABA (GABBR1, GABRB3), and acetylcholinergic receptors (CHRNA1, CHRNB1, CHRM3) also display consistent expression across both systems. F) AMPA receptors (AMPARs) are predominantly expressed by both progenitors and neuronal-like cells in primary tumors and in so-HOTT. Each cell was assigned an NTR identity at the receptor family level based on a two-step gating strategy that prioritizes both enrichment and specificity (see Methods). Stacked bar plots show the distribution of NTR identities across progenitor and neuronal-like populations in primary tumors and in so-HOTT. Cells that did not have a clear NTR identity (i.e., not enriched or not specific) were excluded from the analysis. G) Immunostainings (left) and 3D reconstruction (right) showing the co-localization of presynaptic marker SYN1 (magenta) with excitatory postsynaptic marker PSD95 (cyan) in Nestin-positive tumor cells (yellow) from primary tumor sections. Nuclei were stained with DAPI (gray). The representative image was from the core region of tumor SB002. Scale bar for immunostaining = 20 µm; scale bar for 3D reconstruction = 0.5 µm. H) Immunostainings (left) and 3D reconstruction (right) showing the co-localization of presynaptic marker SYN1 (magenta) with excitatory postsynaptic marker PSD95 (cyan) in EGFP-positive tumor cells (yellow) transplanted into so-HOTT. Nuclei were stained with DAPI (gray). The representative image was from the core region of tumor LB5658. Scale bar for immunostaining = 20 µM; scale bar for 3D reconstruction = 1 µm. I) Box-and-whisker plot showing that the fraction of tumor staining colocalizing with PSD95 and SYN1 in primary tumors and in so-HOTT are comparable and consistent with previously reported figures in literature ^1,6^. Pairwise comparisons were performed using two-sided unpaired Wilcoxon rank-sum tests (ns, not significant). J) Box-and-whisker plot showing a positive correlation between AMPA module scores and a calcium activity transcriptomic signature, HD.Calcium ^54^ in primary tumor cells and so-HOTT. Cells in both datasets were binned into Low (Quartile 1), Medium (Quartiles 2-3), or High (Quartile 4) AMPA expression, and HD.Calcium module scores were plotted for the cells belonging to each bin. K) Calcium imaging of so-HOTT GBM cells transduced with EF1⍺-GCaMP8m lentiviruses showing baseline GCaMP8m fluorescence (t = 0, left) that increased in intensity after addition of 100 µM glutamate (t = 95 s, middle) and subsequently abolished by treatment with 1 µM of tetrodotoxin (TTX) (t = 190 s, right). Scale bar = 50 µm. L) Representative calcium traces showing ΔF/Fo (y-axis) over time (x-axis) in so-HOTT at baseline and after glutamate and TTX treatment confirms that the observed calcium peaks are glutamate-induced and action potential (AP)-dependent. M) Donut chart showing that 5.3% of so-HOTT tumor cells (n = 202 across 4 tumors) exhibit calcium peaks that are glutamate-induced and AP-driven (i.e., abolished after 1 µM TTX treatment) N) Schematic showing the experimental design for combining AMPAR/KAR perturbation with GCaMP8m calcium imaging. Usage of a microRNA-adapted short hairpin RNA (shRNAmiR) targeting the AMPAR subunit *GRIA2* or KAR subunit *GRIK3* allowed placement of the knockdown construct at the 3’ UTR of the GCaMP8m calcium indicator and expression from the same EF1⍺ promoter utilized by GCaMP8m, thereby tying GCaMP8m expression with gene silencing. Lentiviruses harboring this “perturb-and-record” construct were generated and transduced into GBM cells transplanted into so-HOTT. so-HOTT organoids were then sliced and subjected to calcium imaging. O) Donut chart of the shRNAmiR distribution of cells with spontaneous calcium peaks tend to occur less in the shGRIA2 and shGRIK3 conditions compared to the scrambled condition. P) Forest plots showing that shGRIA2 (p = 0.0252) significantly lowers the odds of spontaneous calcium activity in GBM cells compared to the scrambled control. shGRIK3 also trended towards reduced odds, but the effect was statistically non-significant (p = 0.176). Error bars represent 95% confidence intervals. Total n = 529 cells across 3 tumors. Q) Box-and-whisker plots showing that GRIA2 and GRIK3 silencing reduces the peak calcium response amplitude in GBM cells. Violins show the distribution of maximum ΔF/Fo per cell across n = 3 tumors, grouped by shRNAmiR condition. Boxplots show the median (horizontal black line), interquartile range (IQR; white box), and whiskers extending to 1.5 x QR. Pairwise comparisons were performed using two-sided unpaired Wilcoxon rank-sum tests (**p ≤ 0.01, ***p ≤ 0.001).

To functionally validate whether so-HOTT GBM cells are connected to organoid neurons postsynaptically, we performed AAV1-Cre-based anterograde transsynaptic tracing^51,52^ (**Fig. 2B**). AAV1 viruses harboring an EF1⍺-driven Cre cassette were introduced into a subset of cortical organoids prior to tumor transplantation. Afterwards, both AAV1-infected and uninfected control organoids were transplanted with primary tumor cells pre-infected with lentiviruses encoding a Cre-dependent EF1⍺-double-floxed inverse orientation (DIO)-EGFP reporter followed by a unilateral spacer sequence and a miRFP670 cassette. This particular transgene architecture has been shown to reduce leakiness by preventing Cre-independent recombination events in off-target populations^53^. Unconnected tumor cells in both AAV1-infected and uninfected organoids should express miRFP670 by default, but when tumor cells form synapses with Cre-positive neurons in the AAV1-transduced condition, AAV1-Cre viruses from organoid neurons can spread transsynaptically to induce the expression of the EGFP reporter. Indeed, we observed EGFP+ GBM cells in the vicinity of Cre-positive neurons in AAV1-transduced organoids (**Fig. 2C, SFig. 2A**); this was not observed in organoids that did not receive the AAV1 virus. Staining for connected (EGFP+) and unconnected (miRFP670+) GBM cells further show that approximately 5% of tumor cells express EGFP when Cre is present (**Fig. 2C-D, SFig. 2B**), in line with our initial immunostainings (**Fig. 1J**) and previous reports from primary tumor samples^1,6^. Thus, GBM cells are postsynaptically connected to cortical organoid neurons in so-HOTT.

Previous studies have also shown that GBM cells mainly utilize glutamatergic AMPARs to communicate with presynaptic neurons^1,2,6,7^. To test whether this is preserved in our system, we first compared the expression levels of various NTRs in our primary GBM meta-atlas^24^ and in so-HOTT (**Fig. 2E**). Across both systems, we observed AMPARs to be the NTR family with the highest expression (**Fig. 2E, SFig. 2C**); notably, *GRIA2*, which exists in its calcium-permeable form in gliomas and is validated to be a mediator of activity-dependent GBM proliferation^1^, was the most dominant AMPAR in both primary tumors and so-HOTT. We also observed consistent expression of other NTRs in both primary GBMs and so-HOTT, although to a lesser extent compared to AMPARs. These alternative receptors include the kainate receptors (KARs) GRIK3 and GRIK2, GABA receptors GABBR1 and GABRB3, and acetylcholinergic receptors (CHRNA1, CHRNB1, CHRM3), several of which have been shown to also play a role in neuron-glioma communication^3,4,8,9^.

To determine how these different NTRs are distributed across major GBM cell types, we also assigned each GBM cell an NTR identity based on both enrichment and specific expression of an NTR (Methods). With these assignments, we observed that both progenitor and neuronal-like populations predominantly utilize AMPARs in both primary tumors and so-HOTT (**Fig. 2F**). This is consistent with previous studies showing AMPAR utilization by OPC-like progenitor cells^6^ and NPC/neuron-like populations^2^. Additionally, we tested whether the canonical glutamate transporter in cortical cells, VGLUT1, is present in tumor-transplanted cortical organoids. Our scRNA-seq data confirmed that VGLUT1 (*SLC17A7*) was expressed specifically by excitatory neurons in organoids (**SFig. 2D-F**), indicating that the necessary components for NTS are present in both the presynaptic (organoid) and postsynaptic (tumor) sides of so-HOTT. To verify if excitatory synapses exist at comparable levels between primary tumors and so-HOTT, we performed immunostaining and co-localization analysis of presynaptic and postsynaptic markers with Nestin in primary tumors or EGFP in so-HOTT (**Fig. 2G-H, SFig. 3A-B**). Quantification revealed that, in both systems, the percentage of tumor cells harboring excitatory synapses equates to ∼5-10% (**Fig. 2I**), which is equivalent to figures previously reported in literature^1,6^.

**Figure 3.**
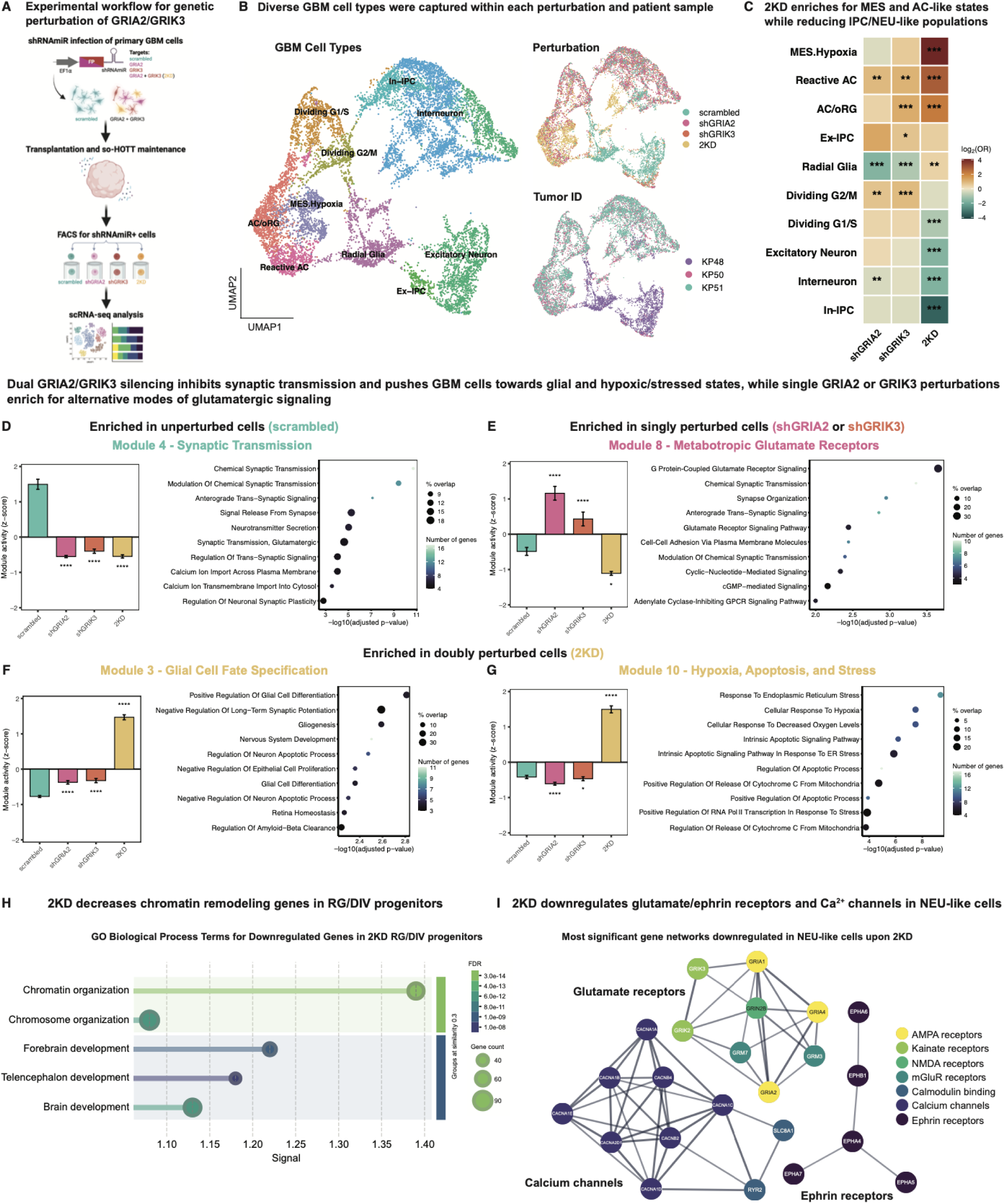
Genetic knockdown of AMPA and kainate receptors alters tumor cell type composition and gene programs A) To genetically perturb AMPAR/KAR subunits in so-HOTT, we transduced primary GBM cells with lentiviruses containing a fluorophore (mCherry or miRFP670) and a scrambled or targeting shRNAmiR sequence that silences *GRIA2* and *GRIK3*. Scrambled cells constituted one cellular pool that was then transplanted into human cortical organoids, while another pool received both GRIA2 and GRIK3 shRNAmiRs, generating four subpopulations (scrambled, shGRIA2, shGRIK3, and double knockdown or 2KD) that were transplanted into a separate set of organoids. After a 2-week period of so-HOTT maintenance, sorting and scRNA-seq was performed on each of the four conditions across n=3 tumors. B) UMAPs of cell type composition after cells were annotated based on a reference atlas of GBM and marker gene analysis show the preservation of diverse GBM cell types after revival from frozen stocks and transplantation into so-HOTT (left). Major cell types were represented across genetic perturbations (right, top) and across tumor IDs (right, bottom). C) Heatmap displaying the log_2_-transformed odds ratio (log_2_OR) for each GBM cell type (rows) across three perturbations (columns) relative to the scrambled control. The diverging color scale represents the log_2_OR derived from a generalized mixed-effects model fit to single-cell binary membership in a given cell type, with tumor ID included as a random intercept to control for inter-tumor variability. Odds ratios were estimated via Wald z-tests on the log-odds contrast between each perturbation and scrambled using estimated marginal means. Positive values indicate that cells are more likely to carry a given annotation in the perturbed condition relative to scrambled, while negative values correspond to depletion. Benjamini-Hochberg (BH) adjusted p-values: *p ≤ 0.05, **p ≤0.01, ***p ≤ 0.001. D) A synaptic transmission module (Module 4) is enriched in scrambled cells but depleted in all three knockdown conditions. Bar plots show the z-scored module activities of selected scNMF modules across the four conditions (left). Asterisks above or below each bar reflect two-sided unpaired Wilcoxon rank-sum tests versus the control. BH-adjusted adjusted p-values: *p ≤ 0.05, **p ≤0.01, ***p ≤ 0.001, ****p ≤ 0.0001). Dot plot shows the top enriched Gene Ontology (GO) Biological Process terms ordered according to level of statistical significance (right). Dot color scale corresponds to the number of genes in Module 4 associated with each term, while dot sizes correspond to the percentage of genes that overlap with the total number of genes for each particular term. E) A module enriched for metabotropic glutamate receptors (Module 8) is enriched in single perturbation conditions. Bar plots (left) and dot plots (right) represent the same information depicted in Figure 3D. F) A glial cell fate specification module (Module 3) is selectively enriched in the 2KD condition. Bar plots (left) and dot plots (right) represent the same information depicted in Figure 3D. G) A hypoxia/apoptosis/stress module (Module 10) is also selectively enriched in 2KD. Bar plots (left) and dot plots (right) represent the same information depicted in Figure 3D. H) Lollipop plots showing the depletion of chromatin organization and neurodevelopment-related genes in 2KD uncommitted RG/DIV progenitors. Enrichment analysis was performed in STRING (version 12.0) using genes downregulated in the RG/DIV maturation group upon 2KD. The y-axis lists significant GO Biological Process terms, while the x-axis shows the signal associated with each term, calculated as a weighted harmonic mean that combines the -log(false discovery rate/FDR) and the observed-to-expected gene occurrence ratio. Color scale represents the FDR, while dot sizes correspond to gene counts for each term. I) STRING interaction network of significantly downregulated genes in NEU-like cells upon 2KD shows depletion of glutamatergic receptors from various subfamilies (AMPA, kainate, NMDA, and mGluR), calcium channels, and ephrin receptors. This high-confidence network was generated by filtering for downregulated genes with interaction scores > 0.90. Enrichment analysis was performed using STRING (version 12.0).

To explore whether AMPAR expression in so-HOTT is correlated with functional measures of synaptic connectivity, we first tested how AMPAR module scores track with a previously published calcium signature (HD.Calcium) in GBM^54^. In this analysis, both primary and so-HOTT GBM cells with high AMPAR scores also exhibited higher HD.Calcium scores (**Fig. 2J**), in line with previous studies showing that glutamatergic signaling promotes calcium activity in tumor cells^2,4,6,7,55^. To explore whether tumor cells in so-HOTT are indeed exhibiting functional calcium activity, we delivered an EF1⍺-GCaMP8m lentivirus into tumor cells prior to organoid transplantation. Calcium imaging of GCaMP8m+ so-HOTT cortical organoid slices showed active baseline signaling, much of which was dependent on putative excitatory synaptic activity as it was activated by glutamate and silenced by tetrodotoxin (TTX) (**Fig. 2K-L**). Overall, we observed that 5.3% of the tumor cells in so-HOTT were responsive to glutamate and had calcium transients that were action potential-dependent (**Fig. 2M**). These results were consistent with our immunostaining, transsynaptic tracing, and existing literature showing excitatory NTS in a similar percentage of GBM cells^1,6^.

Our transcriptional and functional data suggest that so-HOTT effectively mimics the synaptic landscape of primary patient tumors, including utilization of ionotropic glutamatergic receptors (iGluRs). Among AMPAR subunits, *GRIA2* is the most highly expressed across both systems (**Fig. 2E**), as expected from previous reports^1^. Next to AMPARs, we found KARs, particularly *GRIK3*, to be the second most highly expressed subfamily of iGluRs in both primary tumors and so-HOTT (**Fig. 2E, SFig. 2C**). KARs are a class of iGluRs that can carry postsynaptic currents in ligand-gated manner, although they are also known to regulate neurotransmitter release and neuronal circuit maturation and may occasionally act as a non-canonical metabotropic receptor that signals through G proteins^56^. While they are not well studied functionally in the field of cancer neuroscience, previous studies have identified KAR expression in GBM^1^ and a role for GRIK3 in activating RAS/ERK signaling in pediatric pilocytic astrocytomas^57^. We performed immunostaining to validate the primary tumor expression of GRIA2 and GRIK3 alongside a presynaptic marker (VGLUT1), observing co-localization at the junction of tumor and microenvironmental cells (**SFig. 3C-F**). To explore the function of these iGluR subunits in our model, we then designed short hairpin RNAs against *GRIA2* and *GRIK3*, validating that they are effective at knocking down the expression of each target (**SFig. 4A-B**).

**Figure 4.**
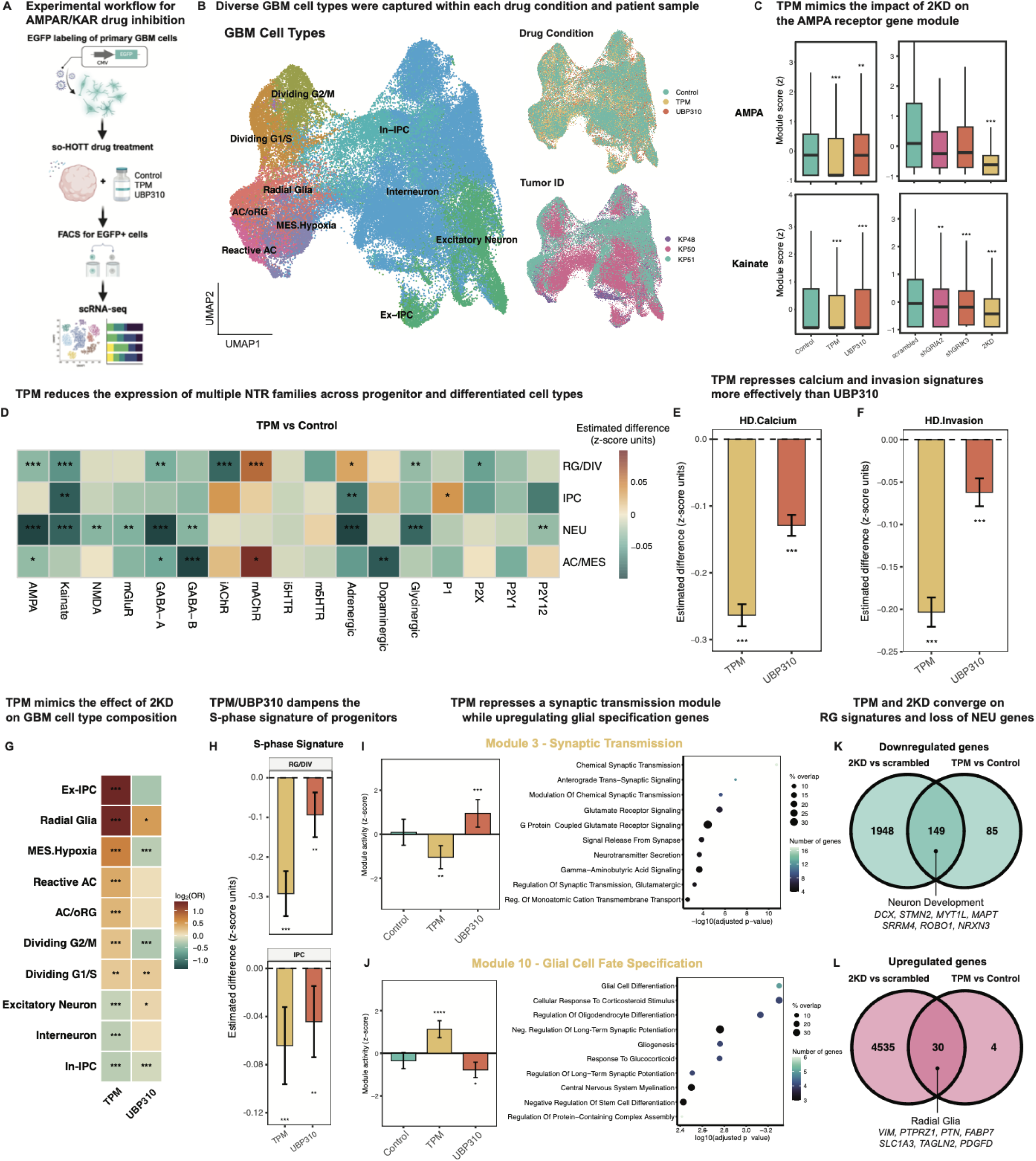
Pharmacological inhibition of glutamatergic synapses modulates GBM cell types and gene programs A) To pharmacologically perturb AMPAR/KAR subunits in so-HOTT, we transduced primary GBM cells with lentiviruses expressing EGFP. Cells were then transplanted into human cortical organoids. After an initial engraftment period of 5 days, so-HOTT organoids were treated with 100 µM topiramate (TPM), a broad AMPAR/KAR inhibitor, or 10 µM UBP310, a kainate-specific inhibitor. DMSO was used as a negative control. Media was exchanged every other day, with drug re-treatments at each media change. After 2-weeks, sorting and scRNA-seq were performed on each of the three drug conditions across n=3 tumors. B) UMAPs of cell type composition after cells were annotated based on a reference atlas of GBM and marker gene analysis show the preservation of diverse GBM cell types after transplantation into so-HOTT (left). Major cell types were represented across drug conditions (right, top) and across tumor IDs (right, bottom). C) Box-and-whisker plots showing the module scores (z-scored across all cells) for AMPA (top) and kainate (bottom) glutamatergic receptors across pharmacological (left) and genetic (right) perturbations show that AMPAR and kainate subunits are downregulated in response to TPM and 2KD conditions. Statistical significance was assessed using a linear mixed-effects model (fixed effect: condition; random intercept: donor). Benjamini-Hochberg (BH) adjusted p-values: *p ≤ 0.05, **p ≤0.01, ***p ≤ 0.001. D) Heatmap of linear mixed-effects model estimates (z-scored module scores) shows that multiple classes of NTRs (x-axis) are downregulated across multiple maturation groups (y-axis) after TPM treatment. Donor information was used as the random intercept to account for inter-tumor variability. The heatmap also shows upregulation of a select group of NTRs, most notably metabotropic acetylcholinergic receptors (mAChRs) in RG/DIV and AC/MES cell types. The diverging color scale represents estimated TPM - Control differences in z-scored module score; color limits are set symmetrically at the 95th percentile of absolute effect sizes across the matrix. BH-adjusted p-values: *p ≤ 0.05, **p ≤0.01, ***p ≤ 0.001. E) Forest bar plot showing the downregulation of a calcium activity signature (HD.Calcium) across all cells in TPM and UBP310 relative to Control. A stronger effect was observed with TPM treatment. The y-axis represents the estimated difference in z-scored module scores between each drug condition and Control, derived from a linear mixed-effects model. Error bars represent 95% confidence intervals. BH-adjusted p-values: ***p ≤ 0.001. F) Forest bar plot showing the downregulation of an invasion signature (HD.Invasion) across all cells in TPM and UBP310 relative to Control. Statistical analysis was done as in Figure 4E. G) Heatmap displaying the log_2_-transformed odds ratio (log_2_OR) for each GBM cell type (rows) across TPM and UBP310 (columns). The diverging color scale represents the log_2_OR derived from a generalized mixed-effects model fit to single-cell binary membership in a given cell type, with tumor ID included as a random intercept to control for inter-tumor variability. Odds ratios were estimated via Wald z-tests on the log-odds contrast between each perturbation and scrambled using estimated marginal means. Positive values indicate that cells are more likely to carry a given annotation in the drug-treated condition relative to Control, while negative values correspond to depletion. BH-adjusted adjusted p-values: *p ≤ 0.05, **p ≤0.01, ***p ≤ 0.001. H) Forest bar plot showing the downregulation of an S-phase cell cycle signature in drug-treated RG/DIV and IPC progenitors. A stronger effect was observed with TPM treatment. The y-axis represents the estimated difference in z-scored module score between each drug condition and Control, derived from a linear mixed-effects model. Error bars represent 95% confidence intervals. BH-adjusted p-values: ***p ≤ 0.001. I) A synaptic transmission module (Module 3) is selectively depleted with TPM treatment. Bar plots show the z-scored module activities of selected scNMF modules across the three drug conditions (left). Asterisks above or below each bar reflect two-sided unpaired Wilcoxon rank-sum tests versus the control. BH-adjusted adjusted p-values: *p ≤ 0.05, **p ≤0.01, ***p ≤ 0.001, ****p ≤ 0.0001). Dot plot shows top enriched Gene Ontology (GO) Biological Process terms ordered according to level of statistical significance (right). Dot color scale corresponds to the number of genes in Module 3 associated with each term, while dot sizes correspond to the percentage of genes that overlap with the total number of genes for each particular term. J) A glial cell fate specification module (Module 10) is selectively enriched in the TPM condition. Bar plots (left) and dot plots (right) represent the same information depicted in Figure 4I. K) Venn diagram showing the intersection of downregulated genes in the TPM and 2KD perturbations indicates that genetic and pharmacological inhibition of AMPARs/KARs converge on depletion of neural development genes. L) Venn diagram showing the intersection of downregulated genes in the TPM and 2KD perturbations indicates that genetic and pharmacological inhibition of AMPARs/KARs converge on enrichment of classical radial glia markers.

We thus sought to test if the calcium fluxes observed in so-HOTT are mediated by these highly expressed glutamatergic receptors. We engineered a lentivirus that harbors the GCaMP8m calcium indicator and a microRNA-adapted short hairpin RNA (shRNAmiR) targeting *GRIA2* or *GRIK3*, both under the control of an EF1⍺ promoter (**Fig. 2N**). This effectively coupled GCaMP8m expression with gene silencing, simultaneously allowing perturbation and the ability to record calcium activity in the same cell. Calcium imaging revealed that *GRIA2* silencing significantly decreased calcium peak detection in so-HOTT relative to the scrambled control; *GRIK3* silencing also led to reductions but the effect was not statistically significant (**Fig. 2O-P, SFig. 4C-E**). Both *GRIA2* and *GRIK3* silencing, however, significantly reduced the maximum amplitude of calcium responses in so-HOTT GBM cells (**Fig. 2Q**). Although the effects of the *GRIK3* knockdown were collectively less substantial than in the *GRIA2* knockdown, we continued investigating KARs to test whether combinatorial AMPAR/KAR targeting produced effects distinct from single-subunit perturbations.

### Genetic knockdown of AMPA and kainate receptors alters tumor cell type composition and gene programs

Given the known spatial association between neurodevelopmental tumor cell types and the infiltrated brain^4,10–12^ and the role of glutamatergic receptors in the cell type specification of normal neural progenitor cells^13–18^, we hypothesized that GBM cell fates are similarly influenced by glutamatergic input. so-HOTT is an ideal model to test this, given that it promotes NTS formation and allows us to selectively perturb tumor cells and profile their transcriptomic signatures through scRNA-seq. Thus, to determine if glutamatergic signaling through AMPARs/KARs influence GBM cell states, we performed a genetic knockdown experiment using fluorophore-coupled shRNAmiRs targeting the AMPAR *GRIA2* and the KAR *GRIK3* in three primary tumors transplanted into so-HOTT. To test how dual AMPAR/KAR perturbation may impact our selected readouts, a double knockdown (2KD) condition targeting both *GRIA2* and *GRIK3* was also included (**Fig. 3A**). These shRNAmiRs were lentivirally introduced into tumor cells prior to organoid transplantation, thus allowing tumor-specific silencing. After a two-week period of tumor engraftment and proliferation in so-HOTT, GBM cells were then sorted and subjected to scRNA-seq. The results show that we can recapitulate the diversity of GBM cell types across the four perturbations and the 3 tumors (**Fig. 3B, SFig. 5A-C**). DEG analysis revealed numerous upregulated and downregulated genes across cell types (**SFig. 5D-E**), with all three perturbations converging upon impaired synaptic transmission (**SFig. 5F-H**). Cell type composition analysis showed that while single perturbations with shGRIA2 or shGRIK3 alone had modest impact on cell type proportions, 2KD led to significant increases in hypoxic mesenchymal- (MES.Hypoxia), reactive AC-, and AC/oRG-like cell types and a concomitant reduction in Ex-Neu-, interneuron-, and inhibitory intermediate progenitor (In-IPC)-like populations (**Fig. 3C**). Dividing cells were also significantly reduced, consistent with previous animal- and cell-based assays with *GRIA2* or AMPAR perturbations^1,6^. Thus, dual AMPAR/KAR targeting impacts GBM cell type composition to a greater extent than individual AMPAR or KAR silencing.

**Figure 5.**
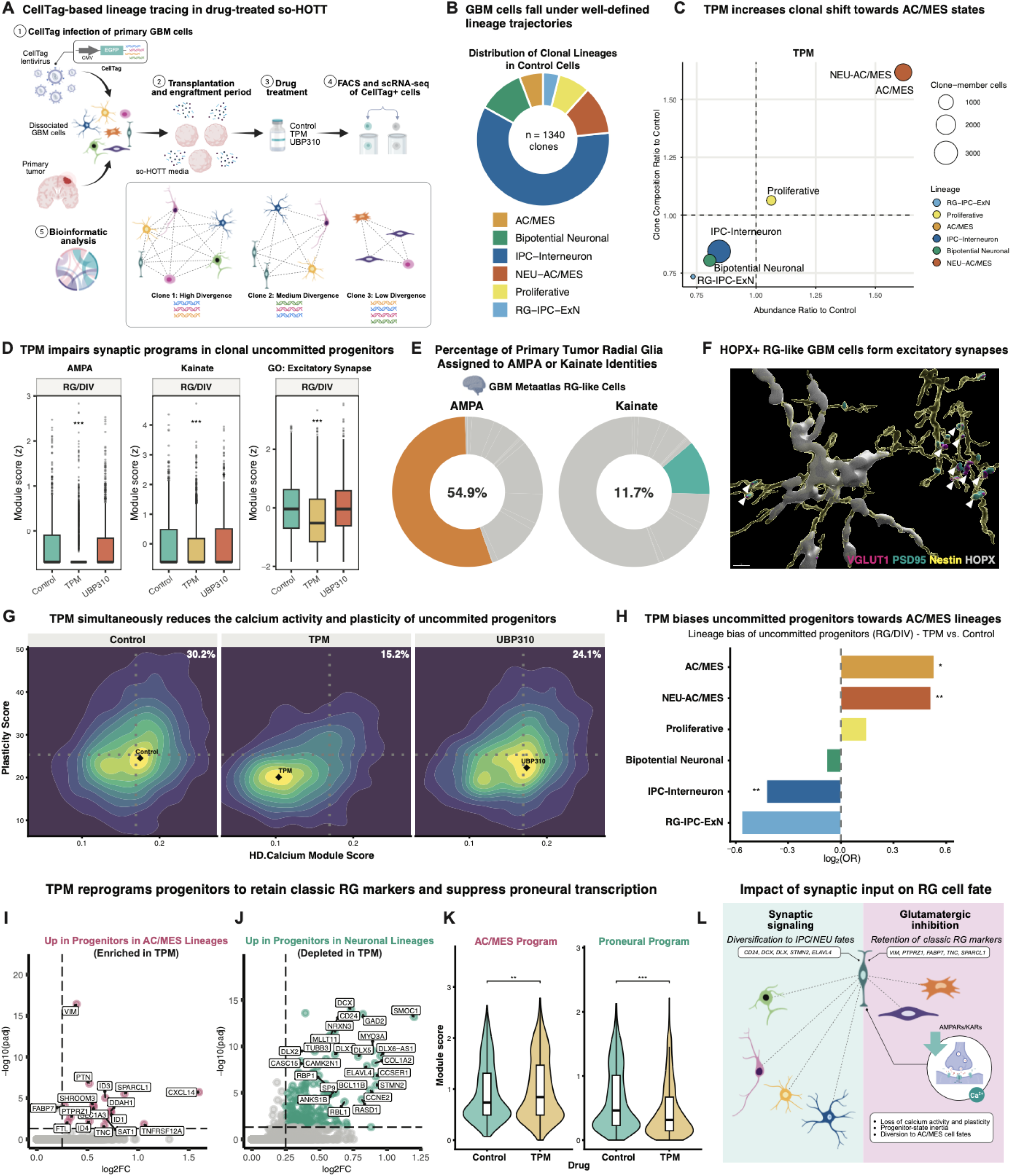
Lineage tracing after AMPAR/KAR blockade reveals clonal reprogramming of radial glia-like tumor progenitors A) To perform lineage tracing, we transduced primary GBM cells from cryopreserved stocks with CellTag lentiviruses, which harbor a library of DNA barcodes that enables identification of tumor cell clones. CellTag+ cells express EGFP, which we used to sort for tumor cells after an initial engraftment period of 5 days and a TPM/UBP310 treatment regimen of 2 weeks. Sorting and scRNA-seq were performed on each of the three drug conditions across n=3 tumors. B) Donut chart showing the baseline distribution of so-HOTT GBM clones from the Control condition across defined lineage trajectories. AC/MES: Astrocytic/Mesenchymal; IPC: Intermediate Progenitor; NEU: Neuronal; RG: Radial Glia, ExN: Excitatory Neuron-Like. C) Clone enrichment (y-axis) versus abundance changes (x-axis) of lineage trajectories in TPM vs. Control indicates enrichment of NEU-AC/MES and AC/MES clones and depletion of neuronal lineages after TPM treatment. Bubble size reflects the number of clone-member cells for each lineage. Clones exceeding 20 cells were excluded to avoid effects of clonal dominance. Colors denote lineage trajectory. D) Box-and-whisker plots showing the module scores (z-scored across all RG/DIV cells) for AMPA (left), Kainate (middle), and a GO: Excitatory Synapse module (right) across drug conditions show that all three gene programs are downregulated in RG/DIV cells after TPM treatment. Statistical significance was assessed using a linear mixed-effects model (fixed effect: condition; random intercept: donor). Benjamini-Hochberg (BH) adjusted p-values: *p ≤ 0.05, **p ≤0.01, ***p ≤ 0.001. E) Donut charts showing the percentage of RG-like primary tumor cells assigned to an AMPA (left) or Kainate (right) identity based on our classification strategy (see Methods). F) 3D reconstruction of primary tumor immunostainings (KP51) showing the co-localization of excitatory presynaptic marker VGLUT1 (magenta) with excitatory postsynaptic marker PSD95 (cyan) within Nestin (NES)-positive tumor cells (yellow) that are positive for the radial glia marker HOPX (gray). White arrows depict colocalizing puncta within HOPX+ NES+ tumor cells. Scale bar: 5 µm. G) Two-dimensional density plots showing the relationship between per-cell HD.Calcium module score and per-cell plasticity scores in uncommitted progenitors. Plots are faceted by Control, TPM, and UBP310 drug conditions. Dotted vertical and horizontal lines represent Control-only median reference values. Black diamonds denote the estimated density peak. The proportion of high-calcium, high-plasticity progenitors, which correspond to cells exceeding both Control median thresholds, are shown in the top right quadrant. H) Waterfall plots showing the significant enrichment of uncommitted progenitors belonging to AC/MES and NEU-AC/MES tracks after TPM treatment. This was accompanied by a decrease in their inclusion in neuronal lineages, particularly the IPC-Interneuron trajectory. The log_2_OR (x-axis) derived from a generalized mixed-effects model fit to single-cell binary membership in a given lineage (y-axis), with tumor ID included as a random intercept to control for inter-tumor variability. Positive values indicate that cells are more likely to carry a given annotation in the perturbed condition relative to scrambled, while negative values correspond to depletion. Benjamini-Hochberg (BH) adjusted p-values: *p ≤ 0.05, **p ≤0.01, ***p ≤ 0.001. I) Right-sided volcano plot showing the upregulation of radial glia markers in uncommitted progenitors that are clonally related to differentiated AC/MES cells. Significance (-log10 of adjusted p-value) is plotted along the y-axis, while the x-axis shows the log2 fold-change. J) Right-sided volcano plot showing the upregulation of proneural genes in uncommitted progenitors that are within neuronal lineages. Significance (-log10 of adjusted p-value) is plotted along the y-axis, while the x-axis shows the log2 fold-change. K) Violin plots showing module scores for the AC/MES program (left) and the Proneural program (right) in uncommitted progenitors in Control vs. TPM conditions. Scores reflect mean normalized expression across signature genes. Inner boxplots show median and interquartile range. Significance brackets indicate pairwise contrasts versus Control from a linear mixed model, with Benjamini-Hochberg (BH) adjusted p-values: **p ≤0.01, ***p ≤ 0.001. L) Model depicting how synaptic signaling through glutamatergic receptors licenses uncommitted progenitors to diversify into neuronal lineages defined by IPC and NEU states by diverging from the RG ground state and inducing the expression of proneural genes. Conversely, inhibition of these receptors leads to loss of calcium activity and constrained plasticity, which is defined by the retention of RG markers and clonal expansion into progenitor-proximal AC/MES states.

To identify latent gene programs that underlying these cell type changes, we performed single-cell non-negative matrix factorization (scNMF) analysis^58^. This revealed 10 gene modules (**SFig. 5I**) that were differentially expressed across perturbations. Notably, Module 4, which included AMPAR/KAR subunit-related genes as well as other genes that facilitate synaptic formation and transmission (e.g., *LRRTM4, DLGAP1*, *DLG2, SYT1, CACNA1E)* were upregulated in scrambled cells, but depleted in all knockdown conditions. This suggested that tumor-intrinsic silencing of *GRIA2* and *GRIK3* not only affects AMPARs/KARs but broadly compromises synaptic signaling (**Fig. 3D**). Interestingly, we also observed the upregulation of Module 8 exclusively in single knockdown conditions. This module is enriched for metabotropic glutamatergic receptors (mGluRs) and associated scaffolding proteins (e.g., *GRM5, GRM7, GRM8*, *HOMER1*), suggesting that single perturbation of GRIA2 or GRIK3, but not 2KD, affords tumor cells some window to adapt or utilize alternative modes of glutamate-based communication (**Fig. 3E**). Lastly, we identified the selective enrichment of glial fate specification (Module 3) and hypoxia/apoptosis/stress genes (Module 10) in 2KD cells. Module 3 notably includes markers for radial glia (e.g., *PTPRZ1, FAM107A, HES1, HOPX*), a developmental cell type that has both neurogenic and gliogenic potential^29^. The expression of cellular stress programs, retention of RG markers, and observed shifts in cell type composition collectively indicate that dual AMPAR/KAR targeting negatively impacts tumor cell health, maintains an RG-like progenitor ground state, and constrains GBM cells towards astrocytic/mesenchymal fates (**Fig. 3F-G**).

Next, we wanted to look at shRNAmiR-induced gene program changes in a cell-type resolved manner. Cell types were first grouped according to maturation levels (**SFig. 5B**), and DEGs were identified within each broader cell type grouping. We looked at which cell types had the largest fraction of DEGs, identifying the greatest shift amongst developmentally uncommitted radial glia (RG), dividing progenitors (DIV), and neuronal-like (NEU-like) tumor cells across shRNAmiR conditions (**SFig. 5D-E**), so we focused further on these. Upon 2KD, uncommitted RG/DIV progenitors displayed downregulation of numerous chromatin remodelling genes, including histone lysine methyltransferases (e.g., *EZH2*, *KMT2A/2C, SETD2*) and histone deacetylases (e.g., HDAC4/9), indicating possible destabilization of epigenetic identities of these progenitors (**Fig. 3H**). On the other hand, NEU-like cells displayed significant reductions not just in the expression of AMPARs/KARs but also glutamatergic NMDA and mGluR receptors (**Fig. 3I**). Additionally, expression of calcium channels and ephrin receptors involved in axon guidance cues were also reduced. This suggests that AMPAR/KAR inhibition not only reduces the fraction of NEU-like cells, but also alters the gene programs of the remaining neuronal-like populations to shift away from iGluR/mGluR usage and calcium-dependent modes of signaling.

### Pharmacological inhibition of glutamatergic synapses modulates cell types and gene programs

To determine if these compositional shifts and functional alterations can be observed in a translationally relevant context, we labeled tumor cells from the same patients as in the previous genetic knockdown experiment with lentiviral EGFP, transplanted them into so-HOTT organoids, and treated them with topiramate (TPM) and UBP310 for two weeks after an initial engraftment period of five days (**Fig. 4A**). We chose these drugs to target the same NTR families targeted in the genetic arm: TPM is a broad AMPAR/KAR inhibitor with clinically-approved applications in epilepsy and migraines^59^, and its inclusion was motivated by our finding that only dual *GRIA2*/*GRIK3* genetic silencing produced strong compositional shifts and gene expression changes. On the other hand, UBP310 is a GRIK1/GRIK3 inhibitor that has not been used in glioma studies but has been shown to reduce epileptic activity by blocking postsynaptic KARs^60^. It was included to specifically isolate the contribution of KARs, which remains largely understudied in GBM. We selected doses based upon the TPM concentration known to inhibit AMPAR/KAR-mediated calcium influx^61^ and a UBP310 concentration that can suppress glutamate-induced currents in *in vitro* systems^62^.

As before, we performed scRNA-seq on the sorted tumor cells, which once again showed preservation of tumor cell type heterogeneity across patients and drug conditions after label-transfer and harmonization with genetic arm annotations (**Fig. 4B, SFig. 6A**). To identify if the drug treatment was functioning on-target, we performed module scoring of NTR families and calculated for the change in NTR module expression across broad classes of cell types, sorted according to previously established maturation groups (**SFig. 5B**). This showed that AMPAR and KAR modules were significantly reduced with TPM treatment across multiple cell types, matching the previous double knockdown condition (**Fig. 4C-D, SFig. 6B**). Meanwhile, UBP310 only reduced AMPAR and KAR modules in neuronal cell types (**SFig. 6B**). Collectively, these results indicate that both TPM and UBP310 were functioning in an on-target manner in our system, though TPM had broader cell type level effects affecting progenitor, NEU-like, and AC/MES populations.

Beyond AMPAR/KAR effects, TPM also resulted in reduced expression of other receptor families (e.g., NMDARs, mGluRs, GABAergic receptors) in NEU-like cells, indicating wider treatment-induced changes in synaptic signaling (**Fig. 4D**). In parallel, we observed upregulation of muscarinic acetylcholinergic receptors (mAChR) in RG/DIV and AC/MES cell types after TPM and UBP310 treatment (**Fig. 4D**). Although individual effect sizes were modest, these shifts were statistically consistent across independent tumors, reflecting reproducible remodeling of NTR identity. Several of these NTRs, including GABAergic and mAChR receptors, have been recently described as mediators of NTS in glioblastoma and diffuse midline gliomas^3,4,8,9^. Together, these shifts in NTR expression suggest the NTR identity may be another axis of variation in GBM, one that is dynamically remodeled in response to synaptic perturbations.

To explore how pharmacological AMPAR/KAR inhibition translate into functional signatures, we focused on the HD.Calcium activity signature^54^ and an invasion signature (HD.Invasion), both of which correspond to processes that are positively correlated with synaptic activity^1,2,4,43^. TPM significantly reduced both HD.Calcium and HD.Invasion signatures across all cell types, while UBP310’s effects were weaker and restricted to fewer cell types (**Fig. 4E-F, SFig. 6C-D)**. These results confirm that pharmacological AMPAR/KAR inhibition leads to meaningful downstream effects on synapse-associated cancer programs.

Having established on-target AMPAR/KAR inhibition, we then explored how these treatments shifted cell type fractions. Composition analysis revealed that TPM, but not UBP310, mimicked the effects of the genetic 2KD condition on cell type proportions. Notably, TPM increased RG-, MES.Hypoxia-, Reactive AC-, AC/oRG-, and Ex-IPC-like cells, while reducing neuronal and In-IPC populations (**Fig. 4G)**. Although the overall numbers of cells annotated as Dividing increased with drug treatment, we also observed that there was a significant decrease in the S-phase signature across progenitor cell types (**Fig. 4H**), which is consistent both with our genetic knockdown experiment and with published reports that inhibiting NTS reduces overall cell division^1,5,6^.

To determine gene programs enriched or depleted in the two drug conditions, we again performed scNMF analysis (**SFig. 6E**). This revealed patterns of module expression that were specifically changed in the setting of TPM treatment. Similar to the 2KD condition in the genetic arm, a synaptic transmission module (Module 3) and a glial cell fate specification module (Module 10) were increased and decreased with TPM treatment, respectively (**Fig. 4I-J**). Cross-application of the genetic arm scNMF modules also revealed similar convergent effects, with 80% of genetic arm modules significantly changing in the same direction in 2KD and TPM conditions (**SFig. 6F**). Lastly, we identified the intersection of downregulated (**Fig. 4K**) and upregulated genes (**Fig. 4L**) in TPM and 2KD cells, which revealed the repression of a neuronal developmental program (e.g., *DCX, STMN2, MYT1L*) and the expression of RG markers (e.g., *VIM, PTPRZ1, FABP7*) in both perturbations. The former supports our previous observation that genetic perturbation of AMPARs/KARs attenuates programs underlying neuronal identity and maturation, while the latter recapitulates the astrocytic/mesenchymal shift previously observed with 2KD.

### Lineage tracing after AMPAR/KAR blockade reveals clonal reprogramming of radial glia-like progenitors

While NTS have been linked to tumor proliferation^1,5,6^, its impact on tumor cell fate trajectories has been unexplored. Previous work has linked NTS to OPC- and NPC-like populations, though other malignant progenitors such as RG-like cells have not yet been identified to have synaptic connections with the microenvironment. Our scRNA-seq of so-HOTT GBM cells after dual AMPAR/KAR inhibition by genetic and pharmacologic means showed shifts in cell type composition as well as gene programs across tumor cell types including progenitors. This led us to hypothesize that NTS may be directly modulating tumor cell fate trajectories by impacting progenitor function. Thus, to evaluate how clonal relationships shift and to explore if progenitor function is altered with AMPAR/KAR inhibition, we coupled lineage tracing analysis with pharmacological perturbation of patient tumor cells in so-HOTT.

so-HOTT is an optimal system to perform lineage tracing because it enables both molecular access to the tumor, scale across tumor cells and patients, and a human-specific synaptic tumor microenvironment. We previously leveraged the CellTagging lineage barcoding system ^63,64^ to understand the lineage architecture of patient tumors^24,44^. We employed the same approach in this study, infecting tumor cells with lentiviruses harboring a barcoded DNA library of ∼19,000 white-listed barcodes prior to organoid transplantation (**Fig. 5A**). Each cell was infected with a target of 3-4 barcodes, leading to an infinitesimally small probability that two tumor cells will randomly receive the same barcode combinations. As a result, cells with significant barcode similarities could be annotated as clones (Methods). Clone calling identified a total of 15,371 clonal members across 4,656 clones spread across 3 tumors, with clones having an average clone size of 3.3 cells (**SFig. 7A**). Importantly, all tumor cell types were represented across clonal cells and clones were equally present in all drug conditions (**SFig. 7B-C**), allowing us to dissect how their relationships changed with drug treatment.

Analysis of clonal populations demonstrated an organization into discrete lineage relationships that recapitulates known neurodevelopmental trajectories (**Fig. 5B, SFig. 7D**). Clones were then assigned to mutually exclusive lineages based on the combination of cell types it contained. Clones composed entirely of dividing (DIV) cells and/or RG-like progenitors, whether homotypic (pure) or heterotypic (mixed), were assigned to the Proliferative lineage, reflecting an uncommitted progenitor state. Clones containing Reactive AC or MES.Hypoxia cells, but without IPC- or NEU-like cells, defined the AC/MES lineage. Clones whose IPC/NEU content was exclusively excitatory, with or without RG/DIV cells, formed the RG-IPC-ExN lineage; conversely, those with exclusively inhibitory members (In-IPC and/or Interneuron), regardless of RG/DIV presence, formed the IPC-Interneuron lineage. Beyond these canonical patterns, clones containing both IPC/NEU-like and AC/MES cells were assigned to the NEU-AC/MES lineage, and clones intermixing excitatory and inhibitory IPC/NEU-like cells, without AC/MES involvement, were designated as Bipotential Neuronal. Importantly, all AC/MES and neuronal sublineages were agnostic to the presence of RG-like progenitors and dividing cells, enabling us to establish clonal links between uncommitted RG/DIV progenitors and more differentiated tumor cell states.

To test how these lineage outcomes were altered by AMPAR/KAR inhibition, we analyzed how TPM and UBP310 treatment alters GBM clonal composition in so-HOTT. TPM enriched for clones belonging to the AC/MES and NEU/AC-MES tracks, in line with previously observed cell type abundance changes (**Fig. 5C**). Conversely, TPM depleted clones belonging to all three neuronal sublineages. On the other hand, UBP310 led to more modest shifts in clonal composition, enriching slightly for NEU-AC/MES clones but to a lesser extent than in the TPM-treated condition (**SFig. 7E-F**).

The striking clonal shifts resulting from TPM treatment prompted us to explore if these lineage outcomes arise from on-target effects of dual AMPAR/KAR inhibition on progenitors. Analysis of clonal uncommitted progenitors revealed that RG-like and dividing cells downregulate AMPAR/KAR and excitatory synaptic gene programs, suggesting direct targeting by TPM (**Fig. 5D**). Importantly, UBP310 had a negligible effect on these modules, mirroring the modest effects that kainate-specific inhibition had on clonal trajectories. We then tested if RG-like cells specifically can harbor excitatory NTS using primary tissue data. Analysis of our GBM meta-atlas shows that 54.9% and 11.7% of primary tumor RGs can be assigned to AMPAR or KAR identities, confirming glutamatergic receptor expression in RGs (**Fig. 5E**). We also immunostained primary GBM tissue to test the presence of these synapses in RG-like tumor cells. Indeed, across 4 tumors, we found excitatory presynaptic (VGLUT1) and postsynaptic (PSD95) puncta colocalizing within Nestin+ cytoplasmic projections of HOPX+ radial glia-like tumor cells (**Fig. 5F, SFig. 7G**). Thus, RG-like GBM cells are synaptically connected and responsive to anti-synaptic drugs.

To further characterize the impact of dual AMPAR/KAR inhibition on these uncommitted progenitors and their clonal fates, we analyzed how TPM impacts the plasticity and calcium activity (HD.Calcium) of these progenitors. To measure the former, we developed a plasticity score, based on a previous approach that measures the transcriptional divergence of a particular clone member from the rest of its clonal partners in a cell-type-agnostic manner^65^. This plasticity metric positively correlated with other annotation-dependent measures of heterogeneity (**SFig. 7H**), including heterotypy (clones harboring mixed cell types) and the Gini-Simpson diversity index, commonly used in ecological studies but has since been applied to measure heterogeneous cell populations^66,67^. As expected, uncommitted progenitors had the highest plasticity scores (**SFig. 7H**), indicating their ability to generate a wide range of cellular identities. The baseline plasticity of each cell type appears to be positively modulated by calcium activity (**SFig. 7I**), prompting us to examine whether inhibition of synaptic signaling restricts progenitor fate potential. Density plots show that TPM treatment significantly shifted uncommitted progenitors, including RG-like GBM cells, towards a low-calcium state that is less plastic (**Fig. 5G, SFig. 8A**), correspondingly depleting the highly plastic, high-calcium pool; in contrast, UBP310 had a comparatively modest effect. This shift was accompanied by reduced clonal diversity: low-plasticity radial glia had clonal partners from a more restricted range of cell types, while high-plasticity ones showed broader fate repertoires (**SFig. 8B-C**). Thus, NTS preserve a progenitor calcium state that enables plasticity; conversely, dual AMPAR/KAR blockade by TPM imposes calcium-dependent constraints on progenitor fate diversification.

To identify the lineages that these progenitors funnel into, we examined their fate bias after treatment with TPM. We observed a significant enrichment of TPM-treated progenitors within astrocytic/mesenchymal lineages (AC/MES and NEU-AC/MES), which coincided with their depletion in various neuronal lineages (**Fig. 5H, SFig. 8D**). Amongst uncommitted progenitors within the AC/MES trajectories, we observed gene signatures that classically mark RG-like cells (**Fig. 5I**), reflecting the progenitor ground state. In contrast, progenitors destined to diversify into neuronal fates downregulated RG markers and instead expressed proneural genes (**Fig. 5J**). This analysis indicates that uncommitted progenitors already express the genes driving their eventual fate, with the AC/MES identity representing a continuation of the RG state, and neuronal identity requiring active transcriptional departure from it. We then treated these gene sets as predictive markers of fate outcomes and tested how their expression changes with AMPAR/KAR inhibition. TPM treatment led to a significant upregulation of the AC/MES program and concomitant downregulation of the neuronal program in uncommitted progenitors, including RG-like cells (**Fig. 5K, SFig. 8E**). Thus, on-target inhibition of AMPAR/KAR activity by TPM leads to the retention of RG genes, shunting them away from neuronal trajectories and instead directing them to clonally expand into progenitor-proximal AC/MES states.

Overall, our data converge on a model (**Fig. 5L**) where glutamatergic signaling in RG-like progenitors enables calcium-dependent cell fate decisions, promoting diversification into neuronal lineages that require progenitors to diverge from their initial RG identity. Conversely, AMPAR/KAR blockade impairs calcium-dependent plasticity, resulting in progenitor retention of classical RG marker genes, which translate into expansion of progenitor-proximal AC/MES cells.

## Discussion

Landmark findings in the field of cancer neuroscience have described the existence of neuron-tumor synapses (NTS), showing across glioma diagnoses that context-specific interference with this synaptic function can decrease tumor division and invasion^1,2,4,6–9,68^. These studies, performed in genetic mouse models, syngeneic transplants, and patient-derived xenograft systems, established tumor innervation as an enabling cancer hallmark^69^. In parallel, single-cell and spatial transcriptomic efforts have shown that GBM is notoriously heterogeneous, with each tumor harboring a unique mix of various cell types with distorted neurodevelopmental-like identities^19–22,24^. This diversity arises through a combination of tumor-intrinsic and -extrinsic mechanisms, including genetics, cell-of-origin effects, hypoxic gradients, immune infiltration, and therapeutic pressures^29^. However, direct investigations on the link between GBM heterogeneity and neuron-tumor synaptic signaling have been limited. This is partly due to the paucity of human-specific models that preserve both NTS and tumor heterogeneity, key features necessary for the simultaneous perturbation of synaptic connections and tracking of GBM cell types and clonal dynamics. To address this, we developed a synapse-optimized human organoid tumor transplantation (so-HOTT) model of GBM. so-HOTT effectively promotes the formation of functional NTS, with 5% of GBM cells synaptically integrating with their neuronal microenvironment. This fraction was consistent across several orthogonal assays, and recapitulated the frequencies that we and others have estimated from primary tumor samples^1,6^. Key molecular features of NTS, including preferential use of glutamatergic receptors and downstream calcium signaling, were also preserved, thus allowing us to faithfully model tumor responses to synaptic perturbations.

Despite having a small fraction of synaptically connected GBM cells, genetic and pharmacological targeting of glutamatergic NTS in so-HOTT led to drastic changes in tumor cell type composition. In particular, dual AMPAR/KAR inhibition shifted tumor cells away from IPC- and NEU-like fates and directed them towards a more homogenous glial identity. These cell type shifts corresponded with changes in the clonal dynamics of RG-like cells, a neurodevelopmental cell type reactivated in GBM^21,70^. These RG-like cells have been previously implicated in tumor proliferation and migration^21,70^; however, whether they are synaptically connected or responsive to NTS inhibition was unknown. Prior neurodevelopmental work have shown that RGs in the ventricular zone (VZ)/subventricular zone (SVZ) express glutamatergic receptors and are responsive to AMPAR/KAR agonists and antagonists^13,15,71^, suggesting that the same may be true for RG-like GBM cells. Here, we discover that RG-like GBM cells are postsynaptic targets of NTS and that these cells alter their plasticity and shift their lineage trajectories in response to glutamatergic inhibition. This finding expands the universe of cell types capable of communicating with the neuronal environment, which was previously thought to be limited to NEU-like, NPC-, and OPC-like cells^2,4,6,12^. Importantly, this places a developmentally uncommitted, multipotent progenitor cell type as a recipient of NTS input and a central node in NTS-dependent cell fate decisions. The involvement of a highly plastic progenitor like RG-like cells explains why perturbing glutamatergic input, despite engaging only 5-10% of tumor cells, can significantly change the types of progeny that emerge.

Apart from deciphering NTS impact on RG dynamics, our results also point to glutamatergic receptor-and calcium-dependent tumor cell plasticity as another key neurodevelopmental feature exploited by GBM cells. Glioma cells have long been known to co-opt normal neurophysiological processes to their advantage: apart from synapses, these cells exploit gap junctions^72–74^ and paracrine factors^45,55,75^ and mimic functional properties of non-malignant cells, including rhythmic calcium oscillations^74,76^, neuronal modes of migration^2^, and adaptive synaptic plasticity^7^. In this study, we find that GBM cells also utilize activity-dependent calcium signaling to regulate cell fate decisions. Mechanistically, AMPAR/KAR blockade attenuated both calcium activity and plasticity of uncommitted progenitors, which became restricted into progenitor-proximal AC/MES states at the expense of IPC/NEU-like progeny. These findings mirror past work in the field of neurodevelopment, which showed that calcium signaling through glutamatergic receptors influence cell fate specification of progenitor cells^13–16,77,78^. Echoing our findings on synapse-dependent tumor fate decisions along a glial-neuronal axis, excitation of neural stem cell progenitors via glutamatergic receptors, voltage-gated calcium channels, and depolarizing GABA signaling have been shown to simultaneously repress glial genes and promote neuronal specification^17,18^. Collectively, these past findings and our results indicate that AMPAR/KAR calcium-dependent signaling is a shared mechanism that can influence the cell fate trajectories of normal and malignant progenitors alike.

What are the translational implications of these findings? Our study essentially reveals that synaptic signaling through glutamatergic receptors enables the plasticity and diversification of RG-like cells into other cell types, thus promoting GBM heterogeneity. Consequently, targeting RG-like cells and their synaptic engagement with their neuronal microenvironment can be a viable therapeutic strategy to attenuate a tumor’s phenotypic plasticity, which was also recently defined as a hallmark of cancer^79^ and is increasingly being appreciated as a catalyst of tumor progression, microenvironmental adaptation, and therapeutic resistance^25,26^. By modulating synaptic input, it may be possible to constrain plastic progenitors to give rise to a more limited repertoire of cell states, thus attenuating intratumoral heterogeneity. While such interventions alone are unlikely to be curative, reducing the complexity of GBM’s cellular landscape represents a significant step toward making these tumors more therapeutically tractable. Our findings provide a rationale for combinatorial regimens that pair NTS targeting with other agents that can target the restricted states that emerge. With the recent discovery of non-glutamatergic modes of neuron-tumor synaptic communication in glioma, future work should systematically profile how the broader spectrum of postsynaptic NTRs beyond AMPARs/KARs can also regulate tumor cell fates. This foundational effort can pave the way for lineage-informed, multi-modal strategies that target synaptic input at multiple nodes to progressively narrow the phenotypic space available to GBM cells.

## DATA AVAILABILITY

Our single-cell RNA sequencing datasets for the HOTT vs. so-HOTT comparison, genetic silencing of AMPARs/KARs, and CellTagging of pharmacologically perturbed so-HOTT tumor cells can be accessed via the UCSC Genome Browser (https://cells-test.gi.ucsc.edu/?ds=gbm-so-hott).

## ACKNOWLEDGMENTS

We thank the members of the Bhaduri Lab at UCLA and Steven Sloan and Tarun Bhatia at Emory University for their insightful advice and feedback on the study. We also thank Jessica Scholes, Jeff Calimlim, and Felicia Codrea at the UCLA Broad Stem Cell Research Center (BSCRC) Flow Cytometry Core for support with cell sorting; Suhua Feng at the UCLA BSCRC Sequencing Core for help with running sequencing; Ken Yamauchi from the UCLA BSCRC Microscopy Core and Soraya Scuderi at the Yale School of Medicine for support and advice on live calcium imaging; Jennifer Achiro and Sylvia Neumann from the Martin Lab at UCLA for providing microscope access and useful information on synaptic assays and drug treatments; Tomas Silva Santisteban at Imaris for advice on image analysis. We also would like to thank Sergey Mareninov, Aung Su, Adrian Murillo, and Fausto Rodriguez at the UCLA Brain Tumor Translational Research Core for enabling tumor sample acquisition. This study was generously funded by support to AB from: NIH NCI P50CA211015 including a Career Enhancement Program Award, The Sontag Foundation (Distinguished Scholar Award), V Scholar Award from The V Foundation, The Uncle Kory Foundation, The American Cancer Society (CSCC-Team-23-980262-01-CSCC), The Margaret Early Medical Research Trust, The Pew Charitable Trusts, The Alexander and Margaret Stewart Trust, the McKnight Neurobiology of Brain Disorders Award, The Rose Hills Foundation, and the Broad Stem Cell Research Center. AM and PRN were funded by the UCLA Eli and Edythe Broad Center of Regenerative Medicine and Stem Cell Research Training Program. BNB is supported by a Canada Graduate Scholarship – Masters from the Canadian Institute of Health Research, a Dorothy May Ladner Memorial Fellowship from the University of British Columbia, a Friedman Award for Scholars in Health from the University of British Columbia and a Canadian Graduate Scholarship – Doctoral from the National Sciences and Engineering Research Council. EF was supported by the David Geffen Scholarship and the UCLA-Caltech Medical Scientist Training Program (T32GM152342). S. Baisiwala was supported by the Tumor Cell Biology (NIH T32 CA-009056) and the Jonsson Comprehensive Cancer Center (JCCC) Cancer Fellowship Awards at UCLA, as well as the Blalock Foundation H&H Lee Resident Research Grant. Additional funding was provided to CVN by the IFER Graduate Fellowship for the Alternative Use of Animals in Science, T32 NS048004, and Predoctoral Fellowship in association with the Training Grant in Neurobehavioral Genetics. RLK was supported by NIH T32GM145388. Lastly, we thank the patients and their families who generously donated and entrusted us with their tumor tissue.

## AUTHOR CONTRIBUTIONS

The study was conceptualized by AM and AB. Experimental designs were made by AM, BNB, and AB. Experiments were performed by AM, BNB, DR, and S. Bollu. EF, S. Baisiwala, WG, RLK, DA, MXL, and TP assisted with experiments, tumor acquisition and processing, and sequencing. Formal analysis was conducted by AM, with support from BNB and DR. EF and PRN assisted with bioinformatic pipeline development. HC supported calcium imaging data processing. Data visualization was performed by AM. Data interpretation was performed by AM and AB. Experimental resources were provided by S. Baisiwala, KSP, and DAN. Project supervision, administration, and funding acquisition were provided by AB. The manuscript was written by AM and AB with input and edits from all authors.

## METHODS

### Stem cell culture and maintenance

UCLA6 human embryonic stem cells (hESCs) were maintained using feeder-free culture conditions, as previously described^37,42^. Cells were cultured on Matrigel-coated 6-well plates in mTeSR Plus medium supplemented with mTeSR Plus Supplement and 1X primocin. Medium was replaced every other day, and cells were passaged once cultures exceeded approximately 75% confluence.

For routine passaging, ReLeSR was added at room temperature for 1 minute and then aspirated. Cultures were incubated at 37°C for an additional 5 minutes before fresh stem cell medium was added to detach colonies into small aggregates, avoiding complete dissociation into single cells. Cell clusters were redistributed onto fresh Matrigel-coated plates at 1:4 or 1:6 split ratios. All cultures were routinely screened for mycoplasma contamination and tested negative. Cell lines were authenticated by short tandem repeat analysis and karyotyping at acquisition from the Broad Stem Cell Core.

### Cortical organoid generation

Cortical organoids were generated as previously described^37,42^. Briefly, at Day 0, hESC cultures exceeding 75% confluence were dissociated by adding 1 mL Accutase per well of a 6-well plate and incubating at 37°C for 5 min. To deactivate the enzyme, 1 mL of Sasai 1 media was then added to each well. Sasai 1 (S1) consists of GMEM supplemented with 20% KnockOut Serum, 0.1 mM β-mercaptoethanol, 1X non-essential amino acids, 1X sodium pyruvate, and 1X primocin. Cells were gently scraped, transferred to 15 mL tubes, centrifuged at 300 x g for 5 min, and resuspended as a single-cell suspension in S1 containing 20 μM Y-27632, 5 μM SB431542, and 3 μM IWR1-endo.

For organoid aggregation, 1 x 10⁶ cells were suspended in 10 mL of S1 containing the three small molecules and distributed evenly into 96-well V-bottom low-attachment plates. Aggregates were left undisturbed for 72 hours. On Day 3, a partial medium change was performed by replacing 50 μL with 100 μL of fresh S1 containing the small molecules to minimize disruption of the newly formed aggregates. Thereafter, complete medium changes were performed every other day. Y-27632 was removed from the medium beginning on Day 7, after the initial aggregation period.

On Day 18, organoids were transferred to low-attachment 6-well plates. From Day 18 to Day 35, organoids were maintained in Sasai 2 (S2), composed of DMEM/F-12 with GlutaMAX, 1X N-2 supplement, 1X lipid concentrate, and 1X primocin. At Day 35, cultures were switched to Sasai 3 (S3), consisting of DMEM/F-12 with GlutaMAX, 1X N-2 supplement, 1X lipid concentrate, 10% fetal bovine serum, 5 μg/mL heparin, 0.5% growth factor-reduced Matrigel, and 1X primocin. Media changes were performed every other day during the S2 and S3 stages. Between Weeks 10-12, organoids were transplanted with tumor cells from fresh dissociations or cryopreserved stocks as described below.

### Tumor transplantation into cortical organoids

Primary tumors were obtained from Ronald Reagan UCLA Medical Center with appropriate consent forms covered by Institutional Review Board IRB numbers 10-000655 and 21-000108. Tumors were cut into smaller pieces and subsequently transferred to a papain solution supplemented with DNase I (Worthington). Papain dissociation was performed over the course of 30-45 minutes at 37°C with vigorous shaking at 5-10 minute intervals. Tumor pieces were further broken down with trituration. For cryopreservation, dissociated tumor cells were pelleted by centrifugation at 300 x g for 5 minutes, resuspended in BamBanker freezing media (Bulldog Bio), stored in freezing containers at -80°C overnight and subsequently transferred to liquid nitrogen tanks for long-term storage. For transplantation of freshly dissociated cells, tumor cells were resuspended in warm Sasai 3 (S3) media after filtration through a 40 µM cell strainer, counted, and subsequently infected with a EGFP lentivirus (SignaGen SL100268) at a concentration of 1 µL per 1 million cells and with polybrene (1:1000) (Millipore Sigma TR-1003-G) for 90 minutes on a rotator at 37°C. For GCaMP8m and genetic perturbation experiments, lentiviruses were delivered at a concentration of 2 µL per 1 million cells, while for CellTagging experiments, the CellTag virus was delivered at a concentration of 50 µL/12 million cells (see below). The same process was applied when infecting cryopreserved cells after thawing in 10 mL of warm media and pelleting. Cells were subsequently washed three times and resuspended in Sasai 3 media at a concentration of 100,000 cells per µL. A total of 1 million tumor cells in a 10 µL volume were then transplanted into cortical organoids using the hanging drop method^37^. The next day, tumor cells were transferred into Sasai 4 media with supplements (S4+) for so-HOTT or S3 media for standard HOTT models. S4+ media consisted of 1X DMEM/F-12 with GlutaMax (Life Technologies 10565-018) with 1X N-2 supplement (Life Technologies 17052-048), 1X CD lipid concentrate (Life Technologies 11905-031), 100 µg/mL primocin (Invivogen NC9141851), 10% Fetal Bovine Serum (HyClone SH30071.03), 10 mg/mL heparin (Sigma H3149), 1% vol/vol Matrigel (BD Biosciences 354230), 1X B-27 supplement (Life Technologies 17054-044), 100 ng/mL soluble NLGN3 (OriGene Technologies TP307955), 10 ng/mL BDNF (Gibco 450-02), and 1 µM each of the following: GSK2879552 (Selleckchem S7796), EPZ-5676 (Selleckchem S7062), N-methyl-D-aspartate (Selleckchem S7072), and Bay K 8644 (Selleckchem S7924). Media changes were conducted every 2-3 days until the point of harvest for immunostainings or single-cell capture.

### Immunostaining

Tumor-transplanted cortical organoids were fixed in 4% paraformaldehyde for 45 minutes at room temperature, while primary tumor pieces were fixed in 4% PFA overnight at 4°C. Fixed specimens were then rinsed with PBS and equilibrated in 30% sucrose in PBS overnight at 4°C. Afterwards, embedding was performed in OCT Embedding Matrix (Fisher Sci 14-373-65) containing OCT (Tissue-Tek, VWR) and 30% sucrose at a 1:1 ratio. Embeddings were then frozen on top of dry ice. The frozen blocks were either transferred to -80°C for long-term storage or cryosectioned as 10-12 µm-thick sections onto glass slides for immunofluorescence staining. To stain the slides, sections were first rinsed with PBS for 15 minutes and then treated with a citrate-based antigen unmasking solution (10 mM sodium citrate, pH 6, Vector Labs) heated to 95°C for 20 minutes. Following antigen unmasking, the sections were permeabilized and blocked with a blocking buffer containing 5% donkey serum, 3% bovine serum albumin, and 0.1% Triton X-100 in PBS for 30 minutes at room temperature. The subsequent primary antibody incubations were performed overnight at 4°C using the following primary antibodies in the blocking buffer: Chicken anti-GFP (1:500, Aves Labs GFP-1020), mouse anti-PSD95 (1:200, Abcam 2723), rabbit anti-SYN1 (1:200, Cell Signaling 5297), chicken anti-Nestin (1:500, Invitrogen PA5-143578), guinea pig anti-VGLUT1 (1:1000, Millipore Sigma AB5905), rabbit anti-GRIA2 (1:200, Proteintech 11994-1-AP), mouse anti-GRIK3 (1:200, Abcam ab233734), rabbit anti-Cre (1:200, Cell Signaling 15036), and rabbit anti-miRFP703 antibody (1:500, Invitrogen PA5 109-200). The primary antibody incubations were followed by three 10-minute PBS washes and secondary antibody incubations with DAPI in the blocking buffer for 2 hours at room temperature. The following secondary antibodies were used. DAPI (1:1000, ThermoScientific 62248), AlexaFluor 488 donkey anti-chicken (1:500, Jackson 703-545-155), AlexaFluor 555 goat anti-guinea pig (1:500, Invitrogen A32849), AlexaFluor 568 donkey anti-mouse (1:500, Invitrogen A32773), AlexaFluor 647 donkey anti-mouse (1:500, Invitrogen A32787), AlexaFluor 647 donkey anti-rabbit (1:500, Invitrogen A31573). Slides were mounted with ProLongTM Gold antifade reagent (Invitrogen) and imaged on a Zeiss LSM 880 confocal laser scanning microscope. Images were thresholded and segmented using a machine-learning based workflow in ilastik (version 1.4.1b23), and segmented images were analyzed for 3-channel colocalization analysis using SynBot^80^. 3D reconstructions were performed in Imaris 11.0.

### Cloning of lentiviral constructs

Three classes of lentiviral constructs were generated for this study: (1) EF1⍺-GCaMP8m for calcium imaging, (2) EF1⍺-GCaMP8m-shRNAmiR for combined calcium imaging and gene knockdown, and (3) EF1⍺-fluorophore (mCherry or miRFP670)-shRNAmiR for gene knockdown with fluorescent cell labeling. The shRNAmiR-containing constructs were built from the pTRIPZ backbone (Addgene 127696), from which the fluorophore and shRNAmiR cassette were amplified by PCR and cloned into pDONR221 via Gateway BP recombination using BP Clonase II (Invitrogen). Target-specific knockdown sequences were introduced by inverse PCR-based site-directed mutagenesis using mutagenic primers to modify the default scrambled antisense guide sequence (5′-ACGTGACACGTTCGGAGAATTA-3′) to either the: GRIK3 antisense sequence: 5′-TTAAAAAAACACAAGATCTGCC-3′; or GRIA2 antisense sequenc: 5′-TTACATTTGAAAAATCACCTGG-3’ For constructs requiring distinct fluorophores, the original fluorophore was replaced with either miRFP670, mCherry, or GCaMP8m by Gibson Assembly (New England Biolabs). The GCaMP8m-only construct was generated separately by Gateway LR recombination using LR Clonase II (Invitrogen) between a pDONR221-GCaMP8m entry clone and the pLEX-305-EF1⍺ lentiviral destination vector. All constructs were packaged into lentiviral particles as described below.

### Lentivirus generation

Lentiviral supernatants were produced by transfecting HEK-293T cells at 90% confluency in 10-cm plates with the envelope plasmid pMD2.G (Addgene #12259), the packaging plasmid psPAX2 (Addgene #12260), and the transfer vector at a 3:9:12 µg ratio using Lipofectamine 2000 (Thermo Fisher). Medium was replaced 24 hours post-transfection with high-glucose DMEM supplemented with GlutaMAX, pyruvate, 10% FBS, and 1X penicillin-streptomycin. Supernatants were collected twice (2 and 3 days post-transfection), pooled, filtered through a 0.45-µm membrane, and concentrated using Lenti-X Concentrator (Takara) at 4°C overnight. Viral pellets were recovered by centrifugation at 1,500 × g for 1 hour at 4°C, resuspended in 1/150th the original volume in PBS, aliquoted, and stored at −80°C until use.

### AAV production and purification

AAV particles were produced in HEK293T cells and extracted using the AAVpro Cell & Sup. Purification Kit Maxi (Takara Bio, Cat. #6676). Extraction solutions were added directly to the culture medium in the 10-cm dish, followed by incubation with the AAV Extraction enhancer at room temperature. The lysate was clarified by centrifugation at 2,000 × g for 10 minutes at 4°C, and the supernatant was concentrated by incubation with Concentrating solution at 4°C for 2–3 hours or overnight. The resulting pellet was recovered by centrifugation, resuspended in Dissolving solution at 1/20th the original supernatant volume, and clarified by a final centrifugation step.

The clarified extract was then purified and concentrated. Residual nucleic acids were digested by incubation with Cryonase Cold-active Nuclease (200 U/mL) at 37°C for 1 hour. Sequential treatment with Precipitator A and Precipitator B followed by centrifugation was used to remove protein and cellular debris. The supernatant was passed through a 0.45-µm Millex-HV filter and transferred to an Amicon Ultra-15 centrifugal filter unit (100 kDa molecular weight cutoff). The AAV solution was buffer-exchanged into Suspension Buffer by five rounds of centrifugation at 2,000 × g at 15°C, with the final concentrate brought to a volume of 800–1,000 µL. The purified AAV was aliquoted into 22 µL single-use aliquots and stored at −80°C until use.

### AAV1-based anterograde transsynaptic tracing

Week 10-12 cortical organoids were infected with AAV1-Cre or received an equivalent volume of PBS-only negative control while singly housed in ultra-low attachment 24-well plates. For the AAV1-Cre condition, organoids were infected with AAV1-EF1⍺-Cre-WPRE viruses at 1:50 dilution, corresponding to 4 µL of virus per organoid in 200 µL media. Organoids were then maintained in a tilted, stationary condition during infection to keep them submerged while minimizing movement, and were incubated for 24 hours at 37°C.

The following day, tumor cells were thawed into warm Sasai 4 medium, pelleted at 300 x g for 5 minutes, resuspended, and counted. Tumor cells were then transduced with EF1α-DIO-EGFP-USS-miRFP670 virus at 2 µL virus per 1 million cells in the presence of polybrene (1:1000). The tumor cell-virus mixture was incubated on a rotator at 37°C for 90 minutes. In parallel, AAV1-Cre-treated and PBS-treated organoids were washed three times with PBS and transferred to warm Sasai 4 medium under shaking conditions. Following viral incubation, tumor cells were washed three times by centrifugation at 300 × g for 5 minutes and resuspended to a final concentration of 1 million cells per 10 µL. Resuspended cells were transplanted into organoids using the hanging drop method. Tumor-transplanted organoids were then kept in ultra-low attachment 6-well plates containing so-HOTT media at 37°C while shaking. Media changes were performed thrice a week, and so-HOTT organoids were then harvested for immunofluorescence staining after 2 weeks.

### GCaMP8m calcium imaging

GBM cells were transduced with a lentiviral EF1α-driven GCaMP8m or EF1α-driven GCaMP8m-shRNAmiR constructs, the latter designed to simultaneously label tumor cells for calcium imaging and perturb AMPAR/KAR signaling. The shRNAmiR cassette encoded either a scrambled control sequence or shRNAmiRs targeting AMPAR/KAR-associated genes, including GRIA2 and GRIK3. Following lentiviral infection, transduced GBM cells were transplanted into human cortical organoids and maintained in so-HOTT maturation media to allow tumor integration and maturation.

For slice preparation, organoids were embedded in 3% low-melting point agarose (Precisionary Instruments) in basal DMEM and sectioned using a Compresstome VF-510-0Z (Precisionary Instruments). Briefly, embedded specimens were inserted into the Compresstome buffer tray filled with ice-cold PBS. Organoids were then sectioned at 300 µm thickness with speed set at 2 and oscillation at 3. Slices were then collected and temporarily maintained in warmed imaging-compatible media until mounting.

For live imaging, organoid slices were mounted onto 35-mm glass-bottom dishes (Cellvis) using 1% low-melting point agarose (Millipore Sigma) in PBS. After immobilization, 300-400 µL of pre-warmed so-HOTT media was gently added to the center of the dish. Mounted slices were then transferred immediately for live fluorescence imaging of GCaMP8m calcium activity.

GCaMP8m fluorescence was recorded from tumor-containing organoid slices using live calcium imaging on a Yokogawa CSU X1 spinning disk confocal microscope mounted on an inverted Zeiss stand, with 488-nm excitation. Spontaneous calcium activity was acquired for 3 minutes at 37°C and 5% CO₂, using a 30-ms exposure time, corresponding to a maximum acquisition rate of approximately 33.3 Hz. For validation of glutamate- and action potential-dependent calcium responses, GCaMP8m activity was recorded sequentially for 3 minutes at baseline, after treatment with 100 µM glutamate, and after subsequent treatment with 1 µM tetrodotoxin (TTX). Regions of interest corresponding to GCaMP8m-positive GBM cells were manually defined using the Graphics function in Zen 3.10 software. Background-corrected fluorescence intensities were exported for each ROI across time and processed using a custom MATLAB pipeline to calculate relative fluorescence changes, expressed as ΔF/F₀.

### Calcium imaging analysis

GCaMP8m fluorescence traces were analyzed using custom MATLAB pipelines. For each imaging file, time-series fluorescence intensities were imported from ROI-level CSV files, using ROI columns corresponding to mean fluorescence intensity measurements. Raw fluorescence traces were baseline-corrected using a rolling median estimate of F0 with a 1000-frame window, corresponding to approximately 30 s at the acquisition rate of 33.33 Hz. Calcium activity was then expressed as ΔF/F_0_, calculated as ΔF/F_0_=(F−F_0_)/F_0_.

Peak detection was performed on the ΔF/F0 traces using MATLAB findpeaks. Peaks were required to exceed a minimum ΔF/F_0_ threshold, satisfy a minimum peak prominence criterion, and be separated by at least 3 s. Peak widths were calculated using the half-prominence reference. For the Baseline-Glutamate-TTX experiments, peaks were detected using a ΔF/F_0_ threshold of 0.04 and minimum peak prominence of 0.03. For the knockdown experiments, peaks were detected using a ΔF/F0 threshold of 0.03 and minimum peak prominence of 0.02. No initial frames were excluded from peak detection.

For the Baseline-Glutamate-TTX experiments, matched ROI traces were analyzed across the three sequential conditions: baseline, 100 μM glutamate, and 1 μM TTX. For each ROI and condition, the pipeline exported ΔF/F_0_ traces, peak counts, mean peak amplitudes, peak amplitudes, and mean peak widths. ROIs were classified according to whether activity was detected in each condition, generating a three-digit binary activity code in the order Baseline-Glutamate-TTX. These classifications were summarized as ROI-level activity pattern tables and pattern count summaries. Cross-condition overlay plots were generated for active ROIs to visualize condition-dependent changes in calcium activity and TTX sensitivity.

For the knockdown experiments, baseline calcium activity was analyzed across control, shGRIA2, and shGRIK3 conditions in three tumors. For each ROI, the pipeline exported ΔF/F_0_ traces and peak summary metrics as above. ROI-level maximum ΔF/F_0_ values were extracted and pooled by tumor and shRNAmiR condition for distributional visualization. ROIs were also classified as active if at least one calcium peak was detected during the recording. For each replicate, the percentage of active ROIs was calculated as the number of ROIs with ≥1 detected peak divided by the total number of ROIs. The MATLAB pipeline generated per-tumor and aggregate summaries, including violin plots of maximum ΔF/F_0_, active ROI summaries, total peak counts by shRNAmiR condition, and pairwise nonparametric comparisons using Wilcoxon rank-sum tests.

Downstream statistical analysis of the shRNAmiR arm was performed in R using a logistic mixed-effects model to account for tumor- and replicate-level structure. The MATLAB-derived replicate-level active ROI table was expanded to one row per ROI, with binary activity status coded as active or inactive. The primary model tested whether shRNAmiR condition altered the probability of ROI activity using the formula:

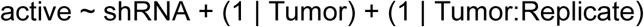

where the shRNAmiR condition was modeled as a fixed effect, with control as the reference level, and random intercepts were included for tumor and replicate nested within the tumor. A null model lacking the shRNAmiR fixed effect was compared with the full model by likelihood ratio test. Fixed-effect estimates were reported as odds ratios with 95% confidence intervals. Pairwise post hoc contrasts between shRNAmiR conditions were performed using estimated marginal means, with Benjamini–Hochberg correction for multiple testing. Model-predicted probabilities of ROI activity were visualized alongside observed replicate-level active ROI percentages, and odds ratios were displayed using forest plots.

### Fluorescence-activated cell sorting (FACS), single-cell capture, and sequencing

Tumor-transplanted cortical organoids were dissociated in papain using the protocol described above. Dissociated cells were suspended in FACS buffer with 2 mM EDTA to prevent cell clumping and DAPI (1:2000) as a live/dead stain. EGFP+ and EGFP- negative live cells (i.e., DAPI-negative) corresponding to tumor and organoid cells, respectively, were then sorted using the BioRad S3e Cell Sorter (HOTT vs so-HOTT comparison) or BD Biosciences FACSAria (genetic and drug arms). EGFP+ (tumor) and EGFP- (organoid) cells across media conditions (HOTT vs. so-HOTT) and tumor regions (Core vs. Periphery) were sorted to characterize the two models. Cells were then captured and processed for particle-templated instant partition sequencing (PIPseq) using the PIPseq V T2 3’ Single Cell RNA Kit (Fluent Biosciences). Tumor cells from the genetic and pharmacological perturbation experiments were sorted based on the fluorophores present in the shRNAmiR lentiviruses (scrambled and shGRIK3: mCherry; shGRIA2: miRFP670; 2KD: mCherry and shGRIA2) and CellTag lentivirus library (See CellTag sections below), respectively. The cells were then captured using the Chromium GEM-X Single Cell 3’ Reagent Kits v4 (10X Genomics). Sequencing libraries were produced according to the manufacturer’s instructions. All libraries were quantified on an Agilent 2100 Bioanalyzer and sequencing was performed on an Illumina NovaSeq 6000 or NovaSeq X Plus (UCLA Sequencing Core).

### CellTag lentiviral transduction, transplantation into so-HOTT, and drug treatment

Freshly dissociated GBM cells were transduced with a CellTag-V1 lentiviral barcode library (Addgene 115643-LVC), which has a complexity of 19, 974 unique barcodes. Briefly, single-cell tumor suspensions were resuspended in Sasai 4 media and incubated with CellTag lentivirus and polybrene (1:1000) at 37°C for 60 minutes with gentle rotation. Lentiviral volume was calculated to achieve a target multiplicity of infection (MOI) of 4 according to this formula:

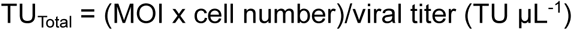

Cells were then washed five times with warm media and resuspended in Sasai 4 using the volume that will yield a concentration of 1 million tumor cells/10 μL. One million CellTag-infected tumor cells were then transplanted into individual Week 10-12 cortical organoids and maintained overnight in a hanging drop configuration as described above. The next day, transplanted organoids were transferred to so-HOTT media and maintained in shaking conditions, with media changes thrice a week. An initial tumor engraftment period of 5 days was observed prior to initiation of drug treatment with topiramate or UBP310. After this period, so-HOTT organoids containing CellTag+ tumor cells were treated with 100 μM topiramate or 10 μM UBP310. A corresponding vehicle control condition treated with an equivalent volume of DMSO was also included. Drug treatments were replenished at every media exchange for 2 weeks until tumor cells were sorted and captured for scRNA-seq as described above.

### CellTag side library generation

To directly associate 10x cell barcodes with CellTag sequences, a targeted side library was generated from DNA extracted from the same 10x Chromium reactions used for transcriptomic profiling. Side library amplification used primers designed to capture the CellTag region downstream of EGFP while appending Illumina-compatible adapters:

> Forward (prtlTruSeqR1_1): ACACTCTTTCCCTACACGACG
>
> Reverse (TruSeqR2-eGFP-3′): GTGACTGGAGTTCAGACGTGTGCTCTTCCGATCTggcatggacgagctgtacaag

The reverse primer annealed downstream of the EGFP coding sequence, ensuring co-amplification of the CellTag cassette and the associated 10x cell barcode. Amplification was performed using the high-fidelity 2X KAPA HiFi HotStart ReadyMix polymerase, followed by indexing PCR to introduce sample indices and full Illumina adapters. Libraries were size-selected and purified using SPRI bead-based cleanup to enrich for the expected fragment size.

### Single-cell RNA sequencing analysis

#### Initial processing, quality control, and cell type annotation

Single-cell RNA sequencing (scRNA-seq) reads were aligned to the GRCh38 human reference genome using PIPseeker software version 02.01.04 (Fluent Biosciences) or 10X Genomics Cell Ranger pipeline (v8.0.0). For the CellTagging experiment, reads were aligned to a modified GRCh38 reference genome that incorporates CellTag UTR and EGFP coding sequences as additional features. Putative doublets were identified using DoubletFinder (v2.0.6). The expected multiplet rate was scaled linearly with cell loading density (0.4% per 1,000 cells, consistent with the 10x Genomics GEM-X platform). DoubletFinder was run using the unadjusted expected doublet count (nExp_poi; pN = 0.25), and all barcodes classified as doublets were excluded prior to downstream analysis. Chromosomal copy number alterations were inferred using inferCNV to distinguish tumor cells from possible non-malignant contaminants. Raw count matrices from CellRanger were loaded, and tumor cells were processed in groups of 2,000 cells per inferCNV run. A common set of 2,000 normal reference cells, downsampled from a normal brain reference object, was appended to each run in subcluster analysis mode using Leiden community detection (PCA embedding; CPM objective; resolution = 0.01; k-nearest neighbors = 80) with denoising and HMM-based copy number state calling enabled (cutoff = 0.1; k_obs_groups = 6). Following each run, the per-cell dendrogram group assignments were extracted across all samples and joined with HMM tumor/normal calls. Cells assigned to dendrogram groups classified as "Tumor" were retained for downstream single-cell analyses.

Using Seurat (v5.2.1), per-sample Seurat objects were then annotated with the corresponding metadata (e.g. Tumor ID, Region, Media, Sample Type, Perturbation, Drug Condition) and merged into a single object after doublet removal and inferCNV. Cells with fewer than 500 unique genes detected (nFeature_RNA < 500) and with >5% mitochondrial gene content were further excluded. Samples were then independently processed through standard Seurat pre-processing: log-normalization, identification of top 2000 variable features, data scaling, and PCA on variable features. The number of principal components retained for each sample was determined empirically using the Marchenko-Pastur eigenvalue threshold^81^. Neighborhood graphs and clusters were generated at a resolution of 0.5 and UMAP embedding were performed on the retained PCs. Objects were then projected onto a pre-integrated GBM meta-atlas^24^ (tumor cells) or developing cortex meta-atlas^82^ (cortical organoid cells from the HOTT vs. so-HOTT experiment) using Seurat’s anchor-based label transfer. Predicted cell identities were used as a guide and coupled with marker gene analysis of Seurat clusters to finalize tumor and organoid cell type annotations. Granular and broad cell type groupings were constructed, the latter based on maturation level of granular cell types.

#### Differential gene expression analysis

Differentially expressed genes (DEGs) were identified using Wilcoxon rank-sum tests (Seurat FindMarkers) and filtered by minimum log₂ fold change of 0.25 and adjusted p-value of ≤ 0.05. DEG discovery was applied at three levels of resolution: globally across all cells, within individual annotated cell types, and within broader maturation groups. To assess overlap across perturbation and drug conditions, Venn diagrams were constructed separately for upregulated and downregulated DEG sets, with all exact overlap regions enumerated and their constituent gene lists carried forward for downstream analysis. Pathway enrichment analysis was performed on each directional DEG set and Venn overlap region using the EnrichR API across selected pathway and gene ontology databases, with visualization of the top enriched terms based on statistical significance and gene overlap. For protein-level interaction analysis, DEG sets were additionally submitted to STRING (version 12.0) and resulting networks were visualized in Cytoscape (version 3.10.4).

#### Cell-cell communication analysis using CellChat

All analyses were performed in R using Seurat (version 5.2.1) and CellChat (version 2.1.2). Briefly, processed and annotated Seurat objects from the PIPseq experiment were loaded for analysis. Samples were subset by region and media condition (so-HOTT vs. HOTT), yielding four main analysis groups: HOTT (Core and Periphery) and so-HOTT (Core and Periphery). Cell-cell communication was inferred using the CellChat package. For each sample group, a CellChat object was initialized (createCellChat()) using normalized gene expression data from the RNA assay and the cell identity metadata as the grouping variable. The CellChat human ligand-receptor interaction database (CellChatDB.human) was used for all analyses. Data were subset to signaling genes, and over-expressed genes and interactions were identified. Communication probabilities between cell groups were calculated (computeCommunProb()) and filtered for robustness. Inferred communication networks and pathway-level results were extracted for downstream analysis. Centrality analysis and signaling role scatter plots were generated to identify dominant sender and receiver populations. Signaling pathway activity was further visualized using bar plots and heatmaps.

#### Module scoring and linear mixed-effects model analysis

Gene modules representing synaptic and functional gene sets were imported from a curated list and applied to the datasets. Briefly, gene module scores were computed from the log-normalized expression matrix for each cell. For each module, all member genes present in the dataset were identified; the score for a given cell was then defined as the arithmetic mean of log-normalized expression values across the detected module genes, divided by the number of genes represented. Modules with no detected member genes were excluded from downstream analyses.

Pre-computed gene module scores were then z-scored globally within each module (across all cells and conditions, pooled) to facilitate cross-module and cross-dataset comparison. To quantify the effect of each experimental condition on module scores, linear mixed-effects models were fit for each module independently using the lme4 package in R, with condition as a fixed effect and donor as a random intercept to account for inter-individual variability. Pairwise contrasts between each experimental condition and the control were extracted via emmeans, and resulting p-values were corrected for multiple comparisons using the Benjamini-Hochberg (BH) procedure. Models were fit at multiple levels of cell-type resolution: pooled across all cells, stratified by granular cell type annotation, and stratified by broader maturation group. Results were visualized using effect-size heatmaps and forest bar plots depicting the mixed-model estimated effect. Box-and-whisker plots also displayed the distribution of per-cell z-scored module scores across conditions.

#### Assignment of neurotransmitter receptor (NTR) identity

A unique neurotransmitter receptor (NTR) identity was assigned to each tumor cell using a two-gate classification strategy applied to pre-computed module scores derived from a curated set of NT receptor gene modules. The first gate, which corresponds to intra-cell specificity, evaluated the relative margin between a cell’s top-scoring and second-scoring NT modules, defined as (score₁ − score₂) / score₁, with a threshold of >0.20 required to pass. The second gate, which corresponds to inter-cell magnitude, z-scored each module’s scores across all cells in the dataset and required the top module’s z-score to exceed 0.50. Cells were then assigned to one of four mutually exclusive classes in priority order: cells in which no module reached the z-score threshold were labeled *Low-NT*; cells passing the first but failing the second gate were labeled *Unassigned*; cells failing the first gate but with at least one module above the z-score threshold were labeled *Ambiguous*; and cells passing both gates were assigned the identity of their top-scoring NT module. This classification was applied independently across experimental arms, with the resulting NTR identity stored as a cell-level metadata annotation for downstream compositional and trajectory analyses.

#### Single-Cell Non-Negative Matrix Factorization (scNMF)

Gene expression programs were identified by applying single-cell non-negative matrix factorization (scNMF) to tumor cells. Cells were subsetted into perturbation or drug conditions. To ensure computational tractability while preserving representation across conditions, up to 3,000 cells were randomly sampled per perturbation group. Log-normalized expression values from the RNA assay were used as input. Highly variable genes were identified using the variance-stabilizing transformation (VST) method implemented in Seurat, and the top 2,000 variable features were selected for factorization. Genes with all-zero or all-NA expression across the input cells were excluded prior to fitting. NMF was performed using the Brunet algorithm (Lee & Seung multiplicative updates) as implemented in the R package NMF, decomposing the genes × cells expression matrix into a basis matrix W (genes × modules) and a coefficient matrix H (modules × cells). A factorization rank of k = 10 modules was specified, and 10 independent random initializations were run to maximize solution stability; the run with the best fit was retained (seed = 1234). The top 50 genes per module were identified by ranking genes on their per-module weight in W.

Per-cell module activity scores were derived directly from the H matrix and stored in Seurat cell metadata. Functional annotation of each module was performed by submitting the top 100 highest-weight genes to Enrichr against the following Gene Ontology (GO) databases. Enrichment terms were retained at a Benjamini-Hochberg (BH)-adjusted p-value threshold of 0.05, and the top 10 terms per database were reported.

For each of the 10 scNMF modules, mean per-cell activity scores were computed within each perturbation condition and arranged into a modules × perturbations matrix. This matrix was displayed as a heatmap with row z-scoring applied across perturbation conditions, such that each row reflects relative enrichment or depletion of a module across perturbations. Rows and columns were hierarchically clustered using Ward’s D2 linkage. To produce per-module bar plots whose directionality directly mirrors the row-scaled mean activity heatmap, mean module activity was first computed for each condition. The resulting condition means were then row-z-scored within each module, subtracting the cross-condition mean and dividing by the cross-condition standard deviation, so that bars above zero indicate enrichment relative to the module’s overall average and bars below zero indicate depletion. Error bars represent 95% t-based confidence intervals computed from per-cell activity distributions and transformed using the same row-centering parameters. Statistical significance was assessed with a Kruskal-Wallis test across all four perturbation conditions, followed by two-sided unpaired Wilcoxon rank-sum tests comparing each perturbation to the scrambled control. P-values were adjusted for multiple comparisons using the BH method and are displayed as significance stars above (positive bars) or below (negative bars) each condition (ns: p > 0.05; *: p ≤ 0.05; **: p ≤ 0.01; ***: p ≤ 0.001; ****: p ≤ 0.0001). All analyses were performed in R. Visualization used ggplot2 and pheatmap.

#### Mixed-effects logistic regression for cell type composition

To test whether shRNAmiR-mediated knockdown of GRIA2 (shGRIA2), GRIK3 (shGRIK3), or both simultaneously (2KD) altered the transcriptional composition of GBM tumor cells, we applied generalized linear mixed-effects models (GLMMs) to single-cell annotation data from the cell type classification. For each cell type, a binary outcome was defined at the single-cell level (1 = member of that cell type, 0 = not), and a GLMM was fit for each pairwise comparison against the scrambled control. Tumor of origin was included as a random intercept to account for inter-tumor variability, treating cells from the same tumor as non-independent observations. Models were fit using the lme4 package in R.

Model-adjusted probabilities and odds ratios (ORs) for each perturbation relative to scrambled were extracted from the fitted models, with 95% confidence intervals computed on the log-odds scale and back-transformed. Multiple testing correction was applied across all cell types within each comparison using the Benjamini-Hochberg procedure (FDR ≤ 0.05).

### CellTag-based lineage tracing analysis

#### CellTag library processing and quality control

CellTag sequencing data were processed from main (transcriptomic) and side (CellTag-enriched) libraries across nine samples spanning three GBM cell lines (KP48, KP50, KP51) each treated with Control, TPM, or UBP310. Raw CellTag UMI count matrices were extracted from CellRanger output and converted to long-format tables mapping CellTag identities to cell barcodes. The technical concordance between main and side libraries was evaluated per sample using UMI count correlations, Venn diagram overlap, and presence/absence metrics. Cell barcodes were then quality-filtered by intersecting with a pre-validated list of singlet cells derived from the annotated Seurat object, which had been filtered for minimum gene detection and tumor cell identity using InferCNV.

#### Clone calling and metadata integration

For each sample, CellTag counts from the main and side libraries were unified using a max-count combining rule, and the resulting table was filtered at a stringency threshold requiring each cell to carry a minimum of two distinct CellTags, with an upper bound of 20 to exclude putative multiplets. Tags with fewer than two supporting UMIs were treated as absent via binarization, and pairwise Jaccard similarity between cells was computed based on their binary CellTag profiles. Clonal groups were identified by graph-based connected-component clustering of cell pairs exceeding a Jaccard similarity of 0.7. The resulting clone assignments were integrated back into the Seurat object metadata, annotating each cell with its clone identity under both threshold conditions for use in all downstream transcriptional and plasticity analyses.

#### CellTag quality control visualization and statistics

Following clone assignment, clonal cells were characterized and visualized across multiple quality dimensions using the fully annotated Seurat object. Clone membership was defined using the Min2 threshold, and analysis was restricted to clones with a global membership of 20 cells or fewer to mitigate the influence of potentially artifactual large clones. A standardized panel of visualizations was generated depicting clone composition by cell type and drug condition, clone size distributions, per-cell-type clone membership fractions, and UMAP overlays of clonal identity. Summary statistics, including total clone counts, total clonal cells, and clone size distributions, were computed globally and stratified by tumor of origin and drug condition.

#### Clonal co-enrichment analysis

Clonal co-enrichment analysis was performed using metadata from the CellTag drug-arm Seurat object. Clone membership was defined using Clone_Min2, and analyses were run across selected genetic arm, broad lineage, broad functional, and module-quartile-combined annotation columns. For each annotation, a binary clone-by-cell type matrix was generated indicating whether each clone contained at least one cell from each cell type annotation. Observed co-occurrence was calculated as the number of clones containing each pair of cell types. Expected co-occurrence was estimated under an independence model based on the overall cell type frequencies across cells and the total number of clones. Log2 enrichment was calculated as log2 observed-over-expected co-occurrence, with a small pseudocount added to avoid undefined values. Log2-enrichment matrices were clustered by Euclidean distance and complete-linkage hierarchical clustering. Heatmaps were generated with dendrogram cuts (k = 5) and clusters were annotated based on cell type membership.

#### Lineage classification

Clonal lineages were assigned to each CellTag-defined clone based on the repertoire of cell types present among its members. Each clone was first assessed against four canonical cell type groupings reflecting known neurodevelopmental cell states: an excitatory neuronal group (Radial Glia, Ex-IPC, Excitatory Neuron), a proliferative/progenitor group (AC/oRG, Dividing G1/S, Dividing G2/M), an astrocytic/mesenchymal group (Reactive AC, MES.Hypoxia), and an inhibitory neuronal group (In-IPC, Interneuron). Clones whose members fell exclusively within one of these groups were assigned to the corresponding lineage trajectory. Clones with mixed membership were reclassified using a hierarchical rule set evaluating the co-occurrence of specific cell type combinations. Clones containing IPC- or neuron-like cells from both excitatory and inhibitory lineages in the absence of AC/MES cells were assigned to a Bipotential Neuronal lineage. Clones containing IPC- or neuron-like cells co-occurring with AC/MES cells were assigned to the NEU-AC/MES lineage. Clones containing proliferative or radial glia cells together with excitatory or inhibitory IPC/NEU-like cells, but not AC/MES cells, were absorbed into their respective neuronal lineage tracks. Clones composed exclusively of radial glia and/or dividing cells without any committed IPC- or NEU-like cells were assigned to the Proliferative lineage. This procedure yielded six final lineage trajectories (RG-IPC-ExN, Proliferative, AC/MES, IPC-Interneuron, Bipotential Neuronal, and NEU-AC/MES), which were appended to the Seurat object metadata for all downstream analyses.

#### Drug-associated shifts in lineage composition

To quantify drug-induced shifts in lineage composition, clonal abundance and enrichment were first assessed per treatment condition using bubble plots. For each lineage, abundance was defined as the proportion of cells belonging to that trajectory out of all cells in the condition, and clone enrichment was defined as the ratio of observed to expected clone membership fraction, where the expected fraction equals the lineage’s cellular abundance. Clones exceeding 20 cells were excluded to minimize confounding from clonal dominance. Control-normalized ratio plots were generated by dividing each treatment’s abundance and clone enrichment values by the corresponding Control values to isolate treatment-specific shifts. To formally test these shifts, a mixed-effects logistic regression model was fit for each lineage trajectory with drug condition as a fixed effect and tumor of origin as a random intercept, run separately for TPM vs. Control and UBP310 vs. Control at both the cell and clone level (with each clone assigned to its majority lineage trajectory). Odds ratios were log2-transformed and p-values were corrected using the Benjamini-Hochberg procedure. Results were visualized as log2(OR) heatmaps and waterfall plots.

#### Development and validation of a per-cell plasticity score

To quantify transcriptional divergence within clones in an annotation-agnostic manner, a per-cell plasticity score was computed for all clonal cells (Min2 threshold, maximum clone size of 20) based on their PCA coordinates. For each cell, the plasticity score was defined as the mean Euclidean distance in the top 30 principal component space between that cell and each of its clonemates (i.e., other cells sharing the same CellTag-defined clone of origin). Cells in clones with fewer than two members in PCA space were excluded. This metric captures within-clone transcriptional spread independent of any cell type label, such that a high plasticity score indicates that a cell occupies a transcriptional state substantially different from its clonal siblings, while a low score reflects transcriptional convergence within the clone.

To validate this metric, per-cell plasticity scores were then averaged at the cell type level, and compared to annotation-dependent measures of clone diversity, including any-pair heterotypy rates and Gini-Simpson diversity index. Any-pair heterotypy was computed per clone as a binary indicator of whether a clone contained at least two cells with distinct cell type annotations (i.e., whether any pair of clonemates differed in assigned identity). For each cell type, the percentage of cells residing in heterotypic clones was then calculated as the mean of this binary indicator across all cells of that type, multiplied by 100. Second, the Gini-Simpson index was computed per clone as 1 − Σp²□, where p□ is the proportion of clone members belonging to cell type k. This metric ranges from 0 (all clonemates share the same identity) to 1 (maximum diversity), and captures graded heterotypy rather than a simple binary. The mean Gini-Simpson index per cell type was then derived by averaging clone-level scores across all cells of each type.

#### Joint distribution of calcium activity and transcriptional plasticity across drug conditions

To visualize the joint distribution of calcium signaling activity and transcriptional plasticity across drug conditions, two-dimensional kernel density estimates were computed for each drug condition (Control, TPM, UBP310) using per-cell HD.Calcium module scores on the x-axis and per-cell plasticity scores on the y-axis. Reference thresholds for both axes were fixed to the Control-condition medians, computed once and applied uniformly across all three drug facets, so that drug-induced shifts in distribution could be interpreted against a common baseline. The density peak within each facet was identified as the coordinate of maximum kernel density and marked with a black diamond. The proportion of cells exceeding both Control median thresholds simultaneously (i.e., occupying the high-calcium, high-plasticity quadrant) was calculated per drug condition as the fraction of cells in that facet with values above both reference lines.

---

**Supplementary Figure 1.**
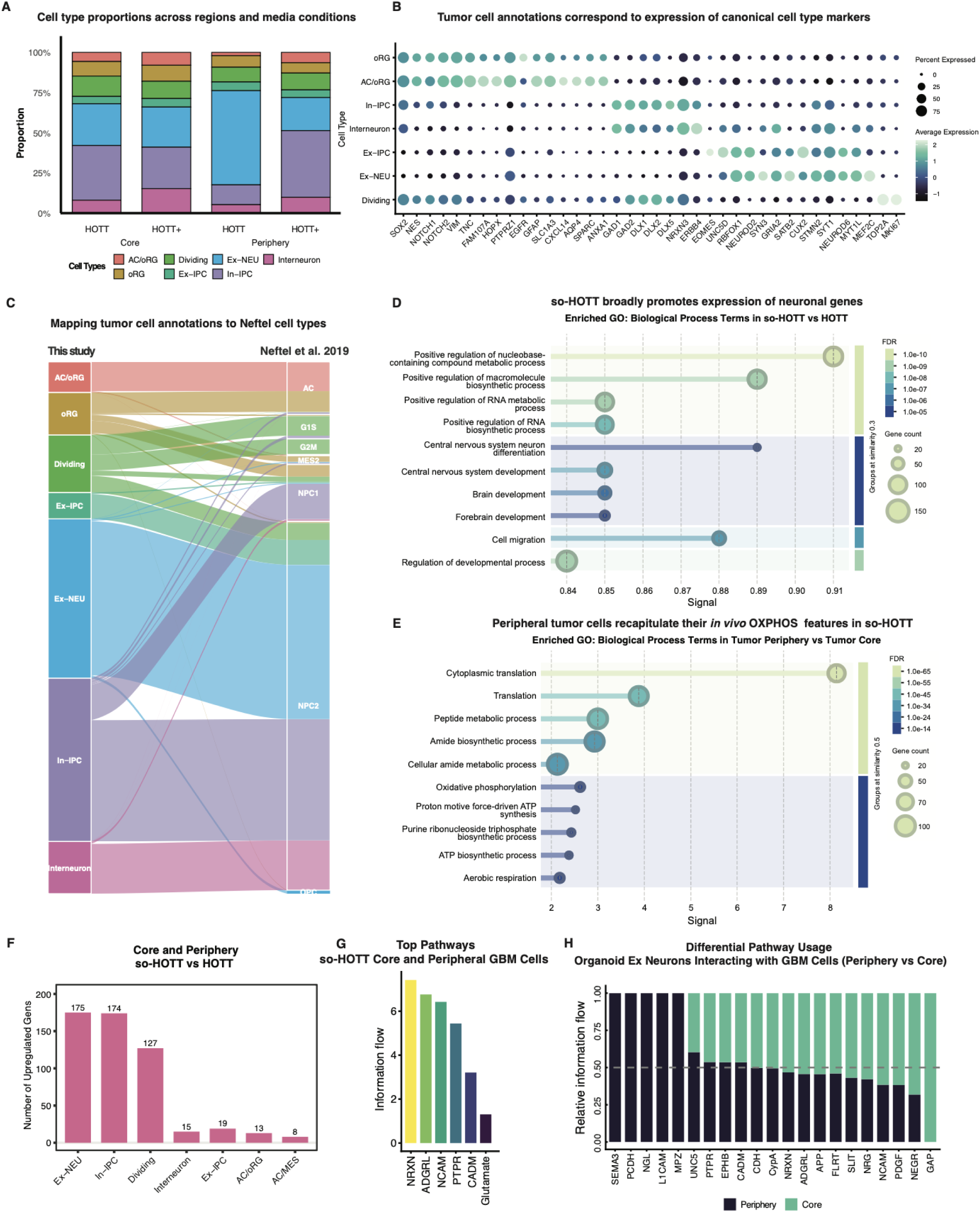
Cell type annotations and differentially expressed genes by media and region. A) Stacked bar plots showing the proportion of each cell type across regions (core vs. periphery) and media conditions (HOTT vs. so-HOTT). B) Dot plot showing marker gene expression (x-axis) across GBM cell types (y-axis) in the HOTT vs. so-HOTT scRNA-seq dataset. Dots are colored by average expression, while dot sizes correspond to the percentage of cells expressing a particular marker within any given cell type. C) Sankey plot showing the correspondence between tumor cell annotations from the HOTT vs. so-HOTT dataset in this study (left) and previously published Neftel cell types (right). D) Lollipop plots showing the enrichment of neuronal, migration, and RNA processing genes in so-HOTT vs HOTT. Enrichment analysis was performed in STRING (version 12.0) using genes upregulated in so-HOTT vs HOTT. The y-axis lists significant GO Biological Process terms, while the x-axis shows the signal associated with each term, calculated as a weighted harmonic mean that combines the -log(false discovery rate/FDR) and the observed-to-expected gene occurrence ratio. Color scale represents the FDR, while dot sizes correspond to gene counts for each term. E) Lollipop plots showing the enrichment of oxidative phosphorylation (OXPHOS) and translation genes in so-HOTT tumor cells from the tumor periphery vs. the core. Enrichment analysis was performed in STRING (version 12.0) using genes upregulated in peripheral so-HOTT tumor cells vs. core so-HOTT tumor cells. The y-axis lists significant GO Biological Process terms, while the x-axis shows the signal associated with each term, calculated as a weighted harmonic mean that combines the -log(false discovery rate/FDR) and the observed-to-expected gene occurrence ratio. Color scale represents the FDR, while dot sizes correspond to gene counts for each term. F) Bar plot showing the number of upregulated genes in so-HOTT relative to HOTT (y-axis) across all cell types (x-axis). Cell types were ranked according to the number of genes per cell type. G) Bar plot showing the information flow (y-axis) for the top CellChat pathways (x-axis) utilized by core and peripheral tumor cells when transplanted into so-HOTT. Information flow is defined as the sum of communication probability among all pairs of cell groups in the network. H) Stacked bar plot showing the relative information flow (y-axis) for the CellChat pathways (x-axis) between peripheral and core tumor cells in so-HOTT. Relative information flow refers to the difference in information flow between two groups.

**Supplementary Figure 2.**
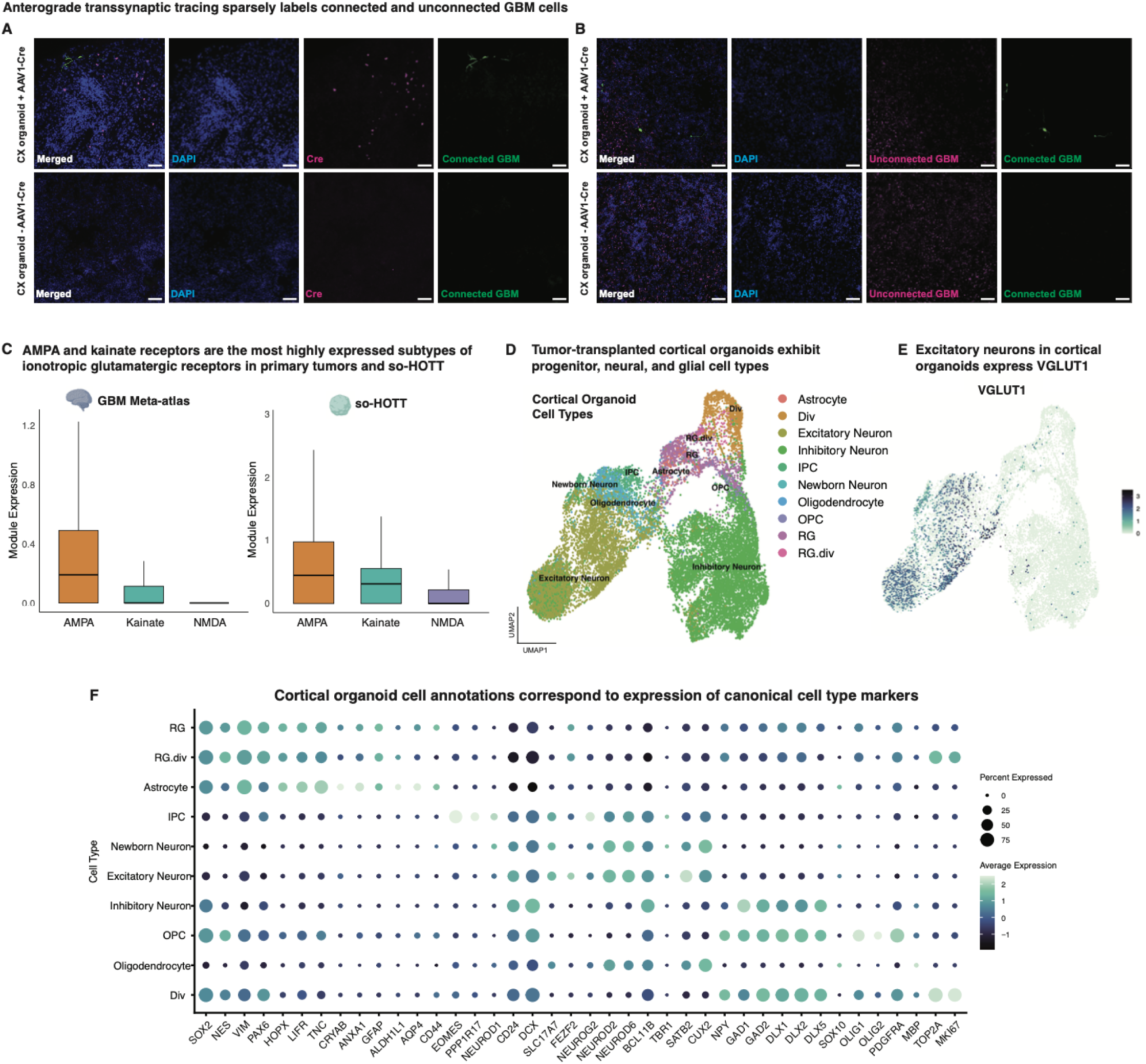
Validation of synaptic features in so-HOTT tumor and organoid cells A) Immunostainings for connected (EGFP+, green) GBM cells in the vicinity of Cre-positive neurons (magenta) in organoids pre-transduced with AAV1-Cre (top row). In organoids without AAV1-Cre, no conversion was observed (bottom row). Scale bar = 50 µm. B) Immunostaining for connected (EGFP+, green) and unconnected (miRFP670+, magenta) GBM cells transplanted into cortical (CX) organoids with (top row) or without AAV1-Cre viruses (bottom row) validates the sparse presence of synaptically connected tumor cells in so-HOTT. Scale bar = 50 µm. C) Box-and-whisker plots showing the module scores (y-axis) for ionotropic glutamatergic receptor subtypes AMPA, kainate, and NMDA receptors (x-axis) in primary tumors (left) and so-HOTT (right). D) UMAP showing the diversity of non-malignant cell types on the cortical organoid side of the assembloid model. E) Feature plot showing the selective expression of VGLUT1 in non-malignant excitatory neurons in cortical organoids. F) Dot plot showing marker gene expression (x-axis) across cortical organoid cell types (y-axis) in the HOTT vs. HOTT+ scRNA-seq dataset. Dots are colored by average expression, while dot sizes correspond to the percentage of cells expressing a particular marker within any given cell type.

**Supplementary Figure 3.**
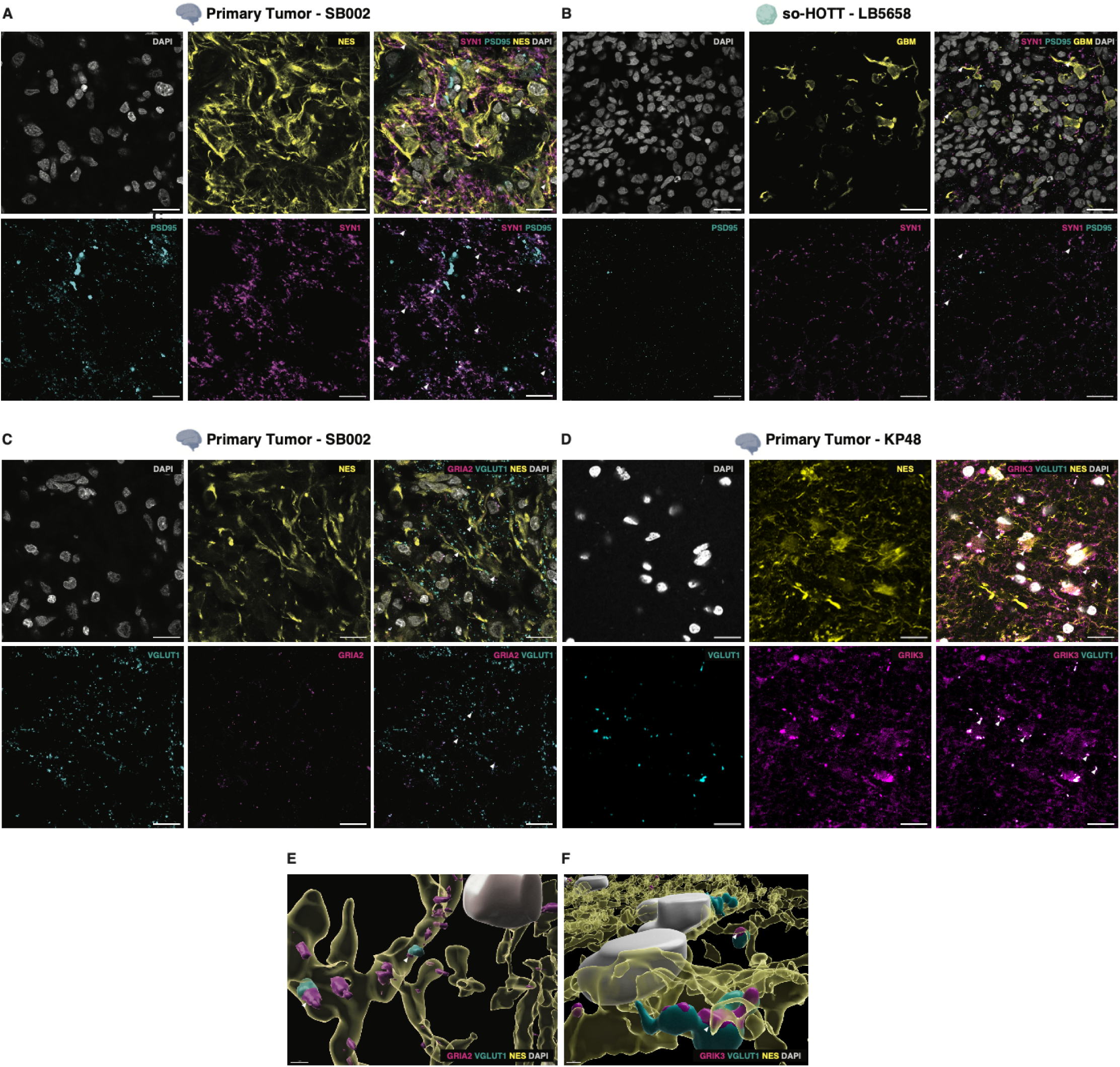
Representative synaptic stainings of primary tumors and so-HOTT A) Immunostaining showing the co-localization (white arrows) of presynaptic marker SYN1 (magenta) with excitatory postsynaptic marker PSD95 (cyan) in Nestin-positive tumor cells (yellow) from primary tumor sections. Nuclei were stained with DAPI (gray). Individual channels (first two columns), as well as two-channel (SYN1 and PSD95) and four-channel merged images (third column), are shown. The representative image was from SB002. Scale bar = 20 µm. B) Immunostaining showing the co-localization (white arrows) of presynaptic marker SYN1 (magenta) with excitatory postsynaptic marker PSD95 (cyan) in EGFP-positive tumor cells (yellow) transplanted into so-HOTT. Nuclei were stained with DAPI (gray). Individual channels (first two columns), as well as two-channel (SYN1 and PSD95) and four-channel merged images (third column), are shown. The representative image shows tumor cells from LB5658. Scale bar = 20 µm. C) Immunostaining showing the co-localization (white arrows) of presynaptic marker VGLUT1 (cyan) with glutamatergic receptor GRIA2 (magenta) in Nestin-positive tumor cells (yellow) from primary tumor sections. Nuclei were stained with DAPI (gray). Individual channels (first two columns), as well as two-channel (VGLUT1 and GRIA2) and four-channel merged images (third column), are shown. The representative image was from SB002. Scale bar = 20 µm. D) Immunostaining showing the co-localization (white arrows) of presynaptic marker VGLUT1 (cyan) with glutamatergic receptor GRIK3 (magenta) in Nestin-positive tumor cells (yellow) from primary tumor sections. Nuclei were stained with DAPI (gray). Individual channels (first two columns), as well as two-channel (VGLUT1 and GRIK3) and four-channel merged images (third column), are shown. The representative image was from KP48. Scale bar = 20 µm. E) 3D reconstruction of VGLUT1+ (cyan), GRIA2+ (magenta) synapses along Nestin-positive tumor cell projections (yellow) in SB002. White arrows depict tumor-localized GRIA2 that is in close proximity to VGLUT1 located outside tumor cell boundaries. Scale bar = 2 µm. F) 3D reconstruction of VGLUT1+ (cyan) GRIK3+ (magenta) synapses along Nestin-positive tumor cell projections (yellow) in SB002. White arrows depict tumor-localized GRIK3 that is in close proximity to VGLUT1 located outside tumor cell boundaries. Scale bar = 1 µm.

**Supplementary Figure 4.**
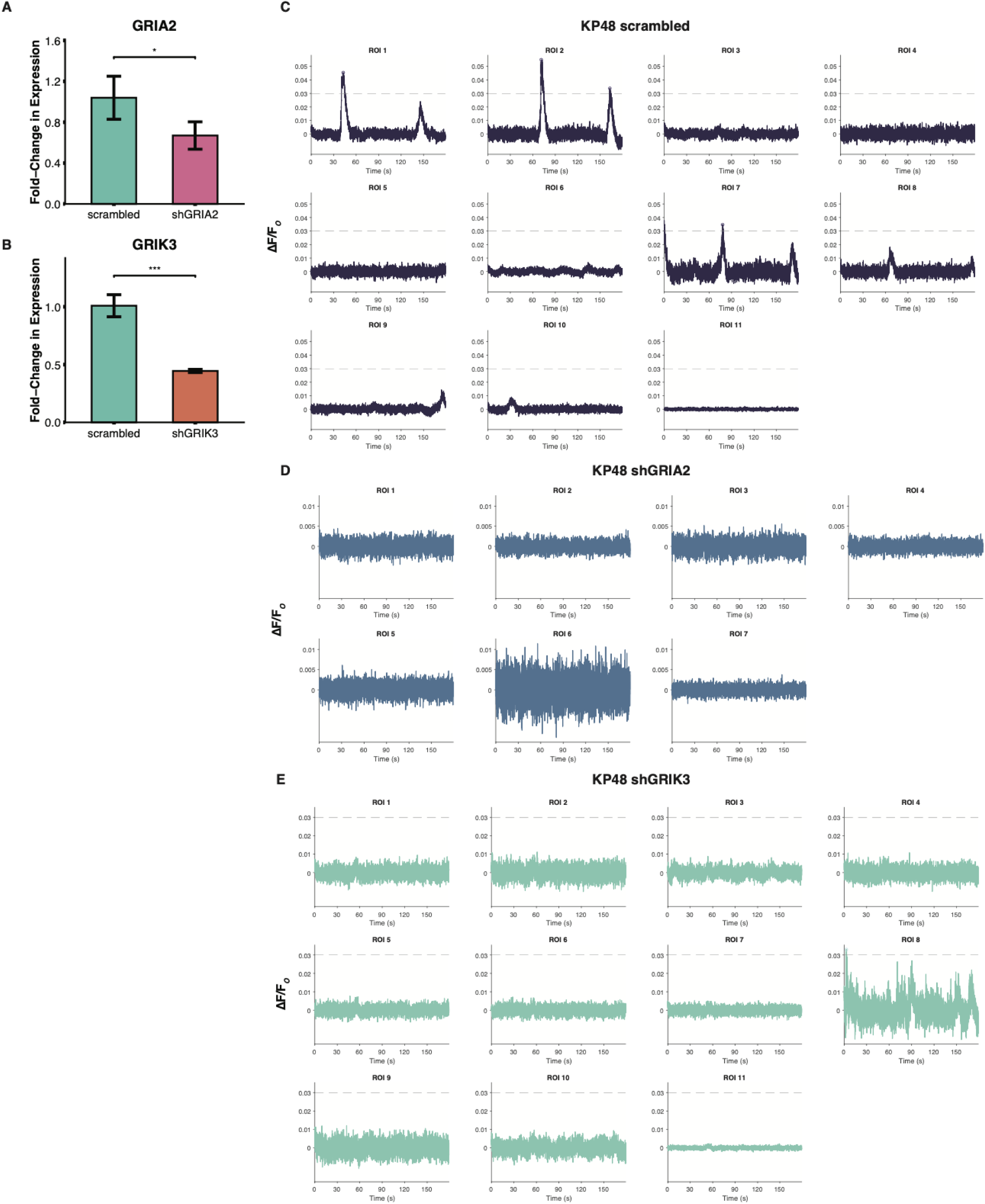
Validation of GRIA2/GRIK3 shRNAmiRs and representative calcium traces A) Bar plot showing quantitative real-time PCR (qPCR) results demonstrating downregulation of *GRIA2* in HK390 glioma stem cells (GSCs) transduced with a lentivirus containing an shRNAmiR targeting GRIA2 (shGRIA2). The statistical significance of the difference between shGRIA2 and the scrambled control was calculated using an unpaired one-tailed t-test (*p ≤ 0.05). B) Bar plot showing quantitative real-time PCR (qPCR) results demonstrating downregulation of *GRIK3* in HK390 glioma stem cells (GSCs) transduced with a lentivirus containing an shRNAmiR targeting GRIK3 (shGRIK3). The statistical significance of the difference between shGRIK3 and the scrambled control was calculated using an unpaired one-tailed t-test (***p ≤ 0.001). C) Representative calcium traces of KP48 so-HOTT GBM cells transduced with the EF1⍺-GCaMP8m-scrambled shRNAmiR construct. Change in initial fluorescence (ΔF/F_O_) over time is plotted for 11 cells from a representative slice. Calcium peaks, which were called using MATLAB’s findpeaks function (Methods), are indicated by calcium elevations that cross the set threshold (dashed line) and are marked by open circles. D) Representative calcium traces of KP48 so-HOTT GBM cells transduced with the EF1⍺-GCaMP8m-shGRIA2 shRNAmiR construct. Change in initial fluorescence (ΔF/F_O_) over time is plotted for 7 cells from a representative slice. No calcium peaks were observed for this slice. E) Representative calcium traces of KP48 so-HOTT GBM cells transduced with the EF1⍺-GCaMP8m-shGRIK3 shRNAmiR construct. Change in initial fluorescence (ΔF/F_O_) over time is plotted for 11 cells from a representative slice. No calcium peaks were observed for this slice.

**Supplementary Figure 5.**
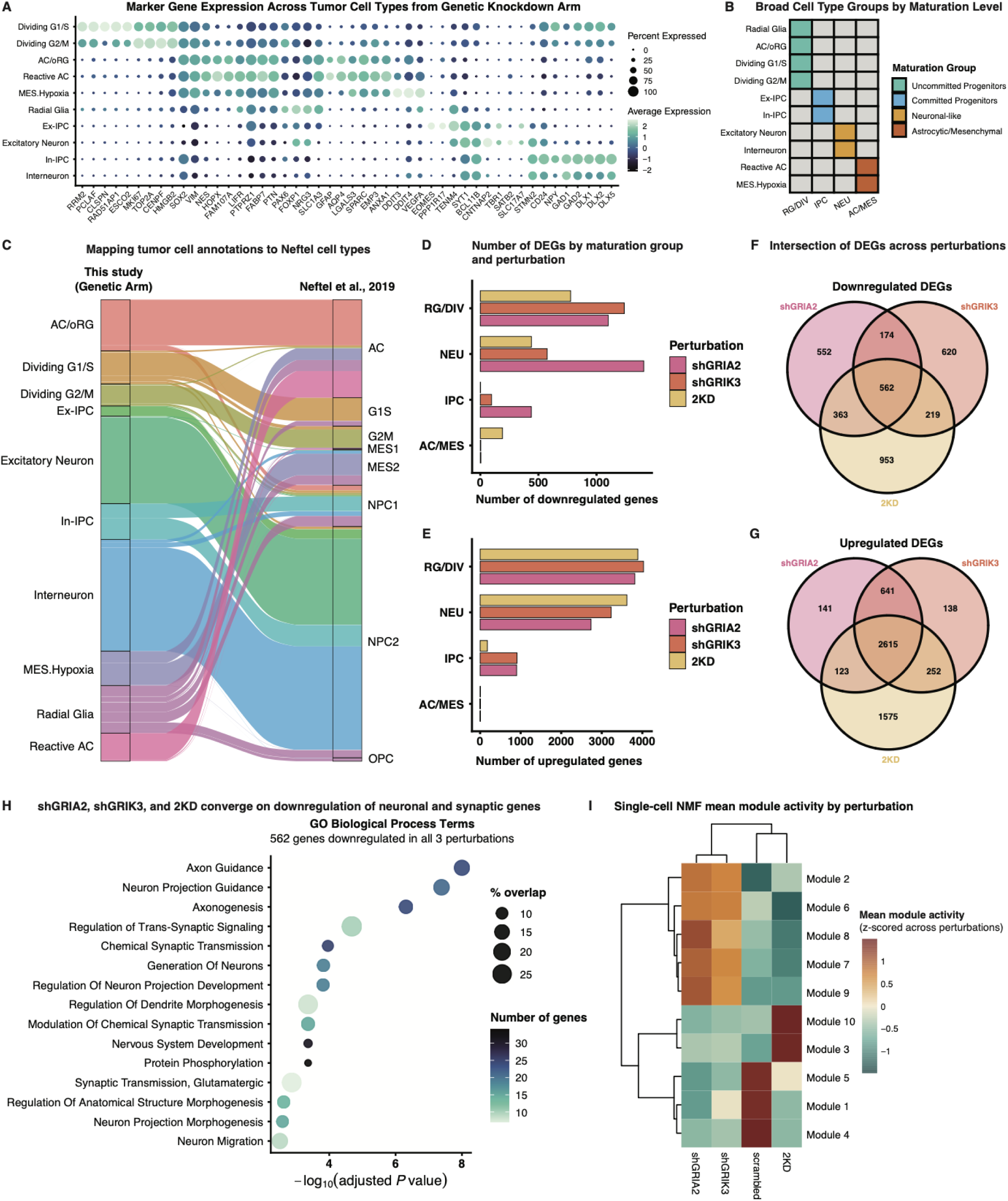
Cell type annotations and gene expression analysis of genetically perturbed GBM cells A) Dot plots showing marker gene expression (x-axis) across GBM cell types (y-axis) in the genetic knockdown scRNA-seq dataset. Dots are colored by average expression, while dot sizes correspond to the percentage of cells expressing a particular marker within any given cell type. B) Binary tile heatmaps showing the aggregation of cell type annotations into broader cell type groups based on maturation level. In certain analyses throughout this paper, radial glia, AC/oRG, and dividing (G1/S and G2/M) have been clustered into uncommitted progenitors (RG/DIV); Ex-IPCs and IN-IPCs into committed progenitors (IPC); Excitatory Neurons and Interneurons into Neuronal-like (NEU); and Reactive AC and MES.Hypoxia into Astrocytic/Mesenchymal (AC/MES). C) Sankey plot showing the correspondence between tumor cell annotations from the genetic knockdown arm of this study (left) and previously published Neftel cell types (right). D) Bar plots showing the number of downregulated genes (x-axis) in each perturbation (color fill) relative to the scrambled control, split across the different maturation groups (y-axis). E) Bar plots showing the number of upregulated genes (x-axis) in each perturbation (color fill) relative to the scrambled control, split across the different maturation groups (y-axis). F) Venn diagram showing the intersection of downregulated genes across the three genetic perturbations. G) Venn diagram showing the intersection of upregulated genes across the three genetic perturbations. H) Dot plots showing enrichment of neuronal and synaptic GO Biological Process terms (y-axis) associated with the 562 genes that are commonly downregulated across the three genetic perturbations. The x-axis shows statistical significance (-log10 of adjusted p-value) of the GO terms listed on the y-axis. Color scale corresponds to the number of genes for each term, while dot size shows the percentage of genes overlapping with the total number of genes for each term. I) Heatmap of mean scNMF module activity scores across perturbation conditions. Each row corresponds to one of the 10 scNMF modules generated for the genetic arm, while each column represents a genetic perturbation. The module activity for each cell is defined as the NMF coefficient from the H matrix (i.e., the weight assigned to that module when reconstructing that cell’s gene expression profile). Mean coefficients were computed per module for each perturbation condition, then row z-scored across the four perturbations, such that the diverging color scale reflects the enrichment (positive values) or depletion (negative values) of each module relative to the cross-condition mean. Hierarchical clustering of rows and columns was performed using Ward’s D2 linkage.

**Supplementary Figure 6.**
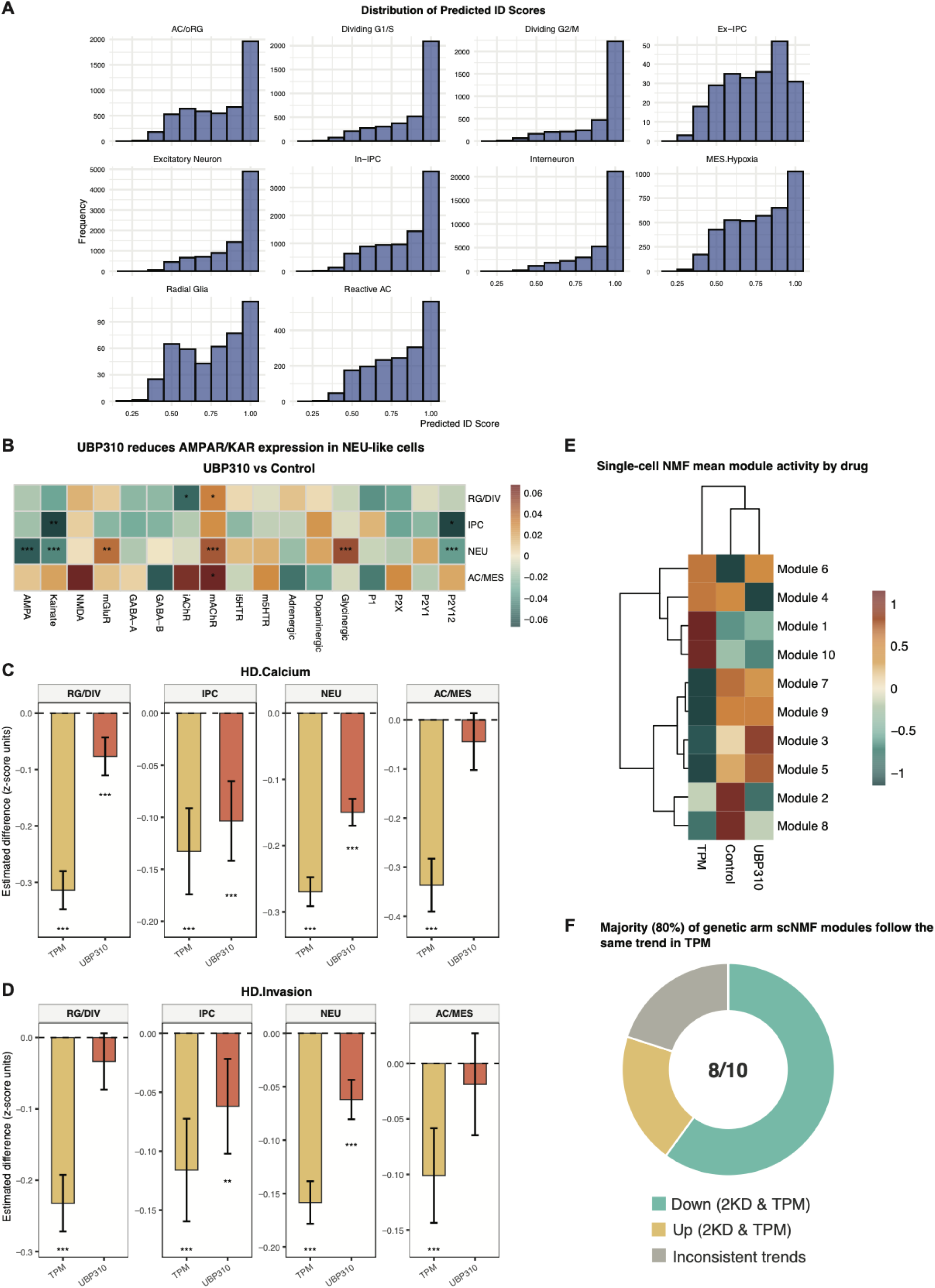
Cell type annotation scores and additional module analysis of drug-treated GBM cells A) Histograms showing the predicted ID scores for each tumor cell type in the pharmacological arm after transferring labels from the genetic arm Seurat object. B) Heatmap of linear mixed-effects model estimates (z-scored module scores) shows changes in NTR module expression across multiple maturation groups (y-axis) after UBP310 treatment. The diverging color scale represents estimated UBP310 - Control differences in z-scored module score; color limits are set symmetrically at the 95th percentile of absolute effect sizes across the matrix. BH-adjusted p-values: *p ≤ 0.05, **p ≤0.01, ***p ≤ 0.001. C) Forest bar plots showing the downregulation of a calcium activity signature (HD.Calcium) across maturation groups in TPM and UBP310 relative to Control. Stronger effects were observed with TPM treatment across broad cell type groups. The y-axis represents the estimated difference in z-scored module scores between each drug condition and Control, derived from a linear mixed-effects model. Error bars represent 95% confidence intervals. BH-adjusted p-values: ***p ≤ 0.001. D) Forest bar plots depicting the same information as Supplementary Figure 7C, but for an invasion signature (HD.Invasion). E) Heatmap of mean scNMF module activity scores across drug conditions. Each row corresponds to one of the 10 scNMF modules generated for the pharmacological arm, while each column represents a drug condition. The module activity for each cell is defined as the NMF coefficient from the H matrix (i.e., the weight assigned to that module when reconstructing that cell’s gene expression profile). Mean coefficients were computed per module for each perturbation condition, then row z-scored across the three conditions, such that the diverging color scale reflects the enrichment (positive values) or depletion (negative values) of each module relative to the cross-condition mean. Hierarchical clustering of rows and columns were performed using Ward’s D2 linkage. F) Donut chart showing percentage of genetic arm scNMF modules that exhibited the same directional trends and statistical significance in the genetic 2KD condition and the pharmacological TPM condition.

**Supplementary Figure 7.**
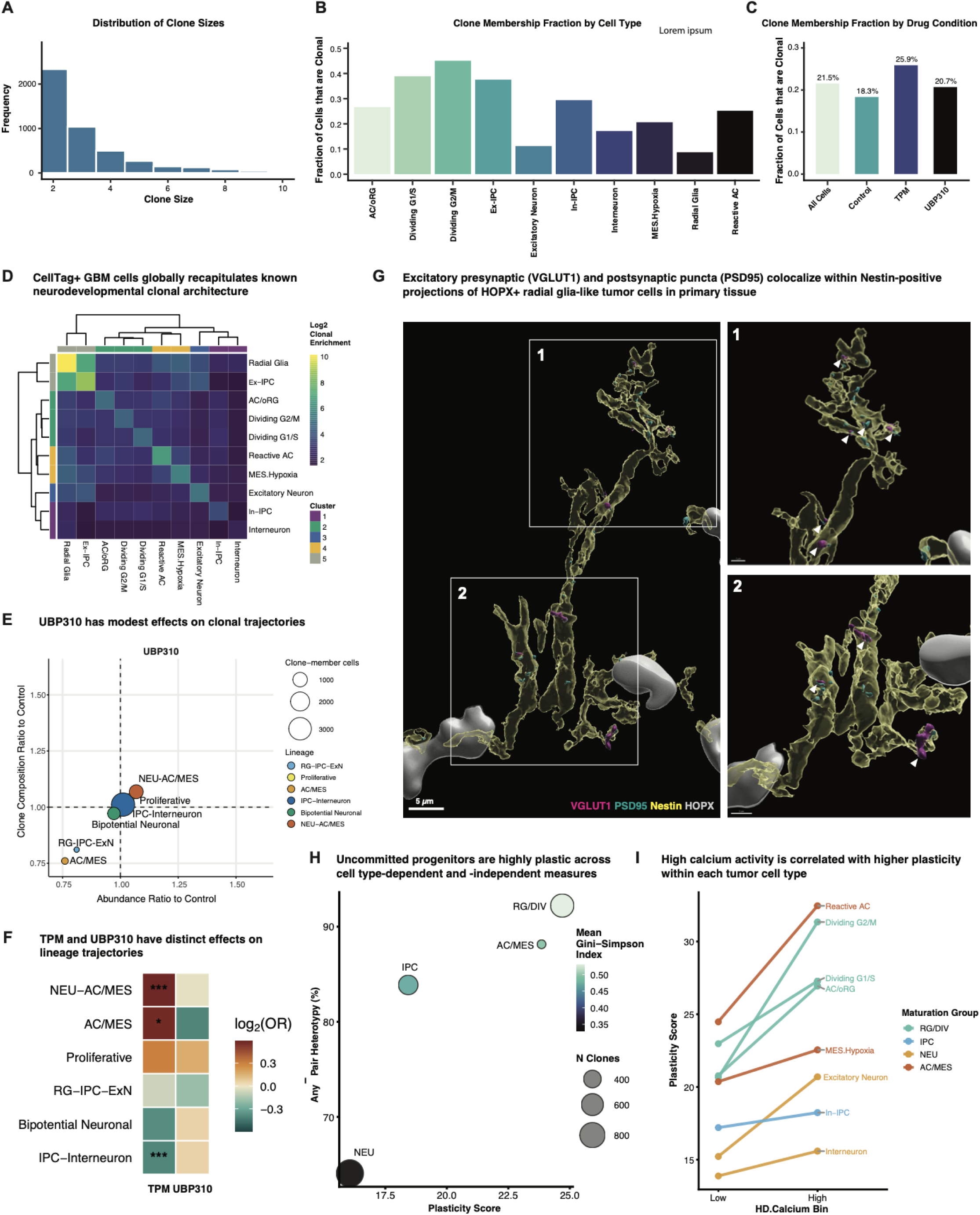
Lineage tracing metrics, radial glia synaptic stains, and plasticity/calcium measures A) Histogram showing the distribution of clone sizes among clones with clone sizes ≤ 20. Each bar represents the number of clones observed at a given clone size. B) Fraction of cells within each cell type that belong to clones (clone size filter ≤ 20 cells per clone). Values were calculated relative to all cells within each lineage state. C) Fraction of cells classified as clone members across treatment conditions (clone size filter ≤ 20 cells per clone). The “All Cells” category summarizes the overall proportion across the full dataset. D) Heatmap showing log2 enrichment of pairwise co-occurrence between annotated cell states within CellTag-defined clones. Each row and column represents a tumor cell type. Enrichment was calculated as the observed number of clones containing both cell types divided by the expected number under an independence model, followed by log2 transformation. Positive values indicate category pairs that co-occur within clones more frequently than expected, whereas negative values indicate reduced co-occurrence. Rows and columns were ordered by complete-linkage hierarchical clustering using Euclidean distance, and colored side annotations indicate dendrogram-defined cluster membership for the selected k value. E) Clone enrichment (y-axis) versus abundance changes (x-axis) of lineage trajectories in UBP310 vs. Control. Bubble size reflects the number of clone-member cells for each lineage. Clones exceeding 20 cells were excluded to avoid the effects of clonal dominance. Colors denote lineage trajectory. F) Heatmap displaying the log_2_-transformed odds ratio (log_2_OR) for each lineage (rows) across TPM and UBP310 (columns). The diverging color scale represents the log_2_OR derived from a generalized mixed-effects model fit to single-cell binary membership in a given lineage, with tumor ID included as a random intercept to control for inter-tumor variability. Positive values indicate that cells are more likely to carry a given annotation in the drug-treated condition relative to Control, while negative values correspond to depletion. BH-adjusted adjusted p-values: *p ≤ 0.05, ***p ≤ 0.001. G) 3D reconstruction of primary tumor immunostainings (SB002) showing the co-localization of excitatory presynaptic marker VGLUT1 (magenta) with excitatory postsynaptic marker PSD95 (cyan) within Nestin-positive tumor cells (yellow) that are positive for the radial glia marker HOPX (gray). Two white boxes demarcate inset regions, with white arrows pointing to colocalizing puncta in RG-like tumor cells. Left panel scale bar: 5 µm; top right panel scale bar: 1 µm; bottom right panel scale bar: 2 µm H) Scatter dot plot showing the relationship between the plasticity score (x-axis) and proportion of heterotypic clones across cell types (y-axis). Color scale depicts the mean Gini-Simpson diversity index of each cell type maturation group, and dot sizes represent the mean clone size for each group. I) Slope plot showing the relationship between calcium activity state and plasticity score across tumor cell types. For each cell type, points represent mean plasticity scores in low versus high HD.Calcium module score bins, with lines connecting the two states. Line and dot color indicates maturation group. Most cell types exhibit increased plasticity scores in the calcium-high state, indicating an association between elevated calcium signaling and greater cellular plasticity.

**Supplementary Figure 8.**
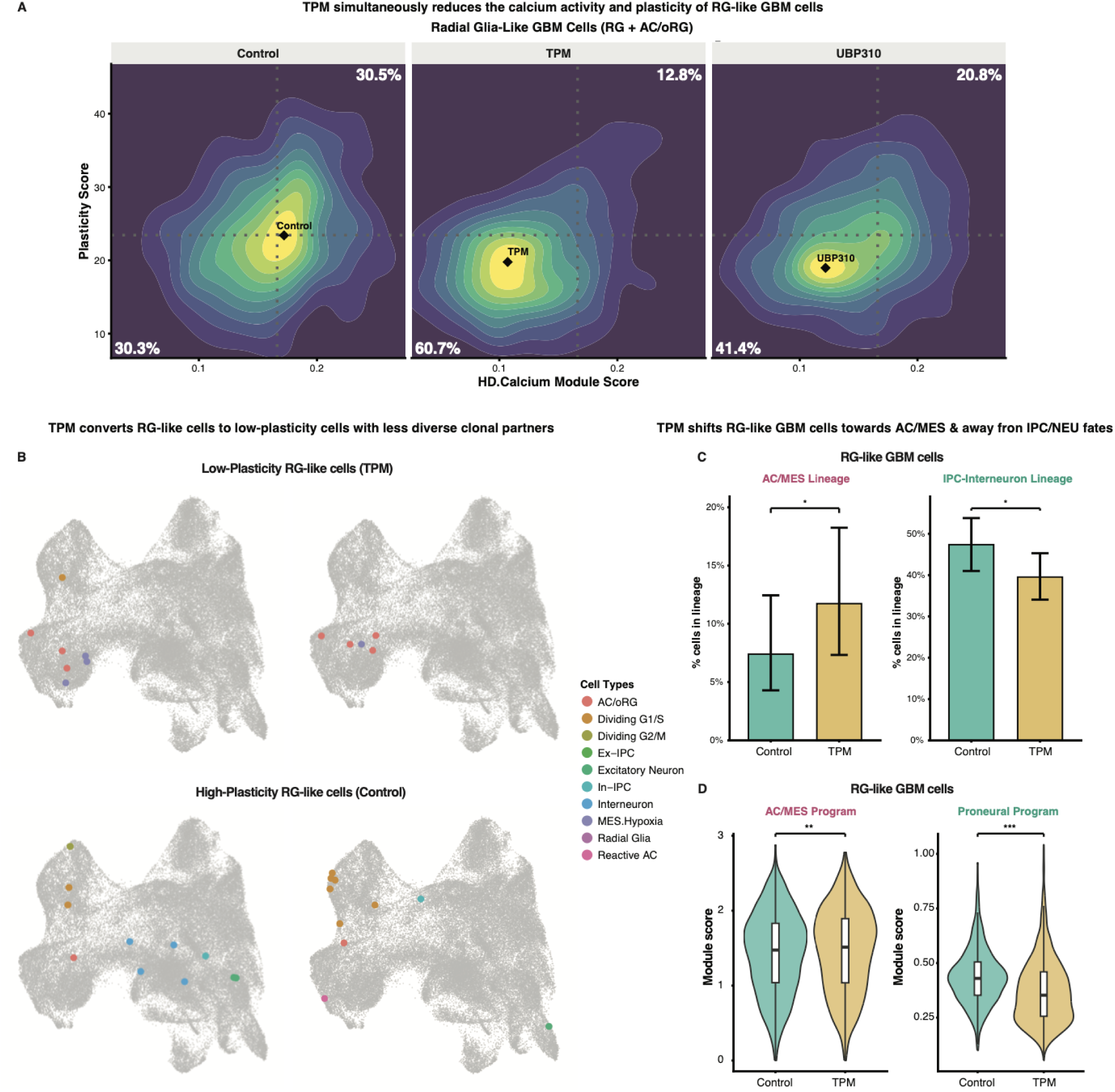
Attenuated plasticity of radial glia-like GBM cells upon AMPAR/KAR blockade A) Two-dimensional density plots showing the relationship between per-cell HD.Calcium module score and per-cell plasticity scores in RG-like GBM cells. Plots are faceted by Control, TPM, and UBP310 drug conditions. Dotted vertical and horizontal lines represent Control-only median reference values. Black diamonds denote the estimated density peak. The proportion of high-calcium, high-plasticity progenitors, which correspond to cells exceeding both Control median thresholds, are shown in the top right quadrant. B) Representative UMAPs showing clonal partners of low- (top) and high-plasticity (bottom) RG-like cells. Clones were stratified by the mean plasticity score of their RG-like cell members. Dot colors denote cell types. C) Bar plots showing the percentage of clonally traced RG-like cells belonging to the AC/MES lineage (left) and IPC-Interneuron lineage (right). Error bars represent 95% confidence intervals. Statistical significance was derived from the pairwise generalized linear mixed model contrasts. *p < 0.05. D) Violin plots showing module scores for the AC/MES program (left) and the Proneural program (right) in uncommitted progenitors in Control vs. TPM conditions. Scores reflect mean normalized expression across signature genes. Inner boxplots show median and interquartile range. Significance brackets indicate pairwise contrasts versus Control from a linear mixed model, with Benjamini-Hochberg (BH) adjusted p-values: **p ≤0.01, ***p ≤ 0.001.

## Notes

### Competing Interest Statement

The authors have declared no competing interest.

